# Dual photoresponsive hydrogel surfaces enabling integrated single-cell analysis of morphology and gene expression

**DOI:** 10.64898/2026.11.18.767535

**Authors:** Xueyang Li, Renjiro Uehara, Koki Mizuno, Sakurako Sato, Arisu Tanji, Kenjiro Tanaka, Nobuo Murase, Shinya Yamahira, Ryuji Kato, Tomoko Yoshino, Akimitsu Okamoto, Satoshi Yamaguchi

## Abstract

Integrated single-cell analysis linking time-resolved cellular phenotypic dynamics with gene expression is essential for understanding cellular heterogeneity but remains technically challenging. Here, we report a dual photoresponsive hydrogel that combines a photoactivatable cell-trapping material with a photodegradable hydrogel, enabling light-directed single-cell capture, time-resolved imaging, and the on-demand release of selected cells for downstream gene expression analysis. This approach allows cells to be arrayed at a high density with uniform spacing, facilitating high-throughput monitoring of morphological dynamics. Cells exhibiting phenotypes of interest can be precisely retrieved by localized hydrogel degradation induced by secondary light irradiation and subsequently subjected to single-cell gene expression analysis. Using this workflow in combination with high-throughput real-time image processing technology, we directly link dynamic morphological features to gene expression and demonstrated the ability of the workflow to examine transcriptional alterations associated with invasive cancer cell behavior.

## Introduction

Recent advances in single-cell analyses have revealed that cellular heterogeneity plays a pivotal role in disease initiation [1], progression [2], and therapeutic response [3]. Accordingly, in fields ranging from fundamental studies of disease mechanisms [4, 5] to drug discovery [6, 7] and diagnostics [8, 9], single-cell analysis has become increasingly important for detection and analysis of previously unknown minor cell populations which are often ignored when averaged across entire major populations [10]. Although genetic sequencing of isolated single cells remains the predominant approach for single-cell analysis, the growing demand to investigate cellular phenotypes, such as morphology [11, 12], motility [13], secretion [14], and cell–cell interactions [15], highlights the increasing need for advanced single-cell analysis techniques that enable direct observation of cellular behaviors [11–15]. Development of an integrated analytical method that directly correlates cellular phenotypes with gene expression profiles will enable obtaining novel insights into diseases, such as cancer and neurological disorders, in which cellular heterogeneity and dynamic gene regulation are key determinants of disease progression and therapeutic outcomes [16, 17].

Image-based single-cell sorting techniques have been developed to meet the requirements of integrated single-cell analysis of cellular phenotypes and gene expression profiles [18–24]. Most of these techniques are based on the use of microstructures such as microwells [18, 19], micropillars [20, 21] and droplets [22], where the cells are confined within narrow microstructures to isolate single cells for high-throughput microscopic imaging. However, these conventional approaches have rarely been applied to phenotypic analysis of adherent cells without specific modifications or adaptations [23], probably because the restricted microenvironments prevent the cells from adopting their native spread morphology, thereby limiting their motility, functionality, and intercellular communication. Additionally, once adhered, such cells are difficult to recover without causing cellular damage [24]. Mechanical or enzymatic detachment often causes cell rupture and loss of genetic material, compromising the accuracy of subsequent single-cell analyses [25]. Therefore, there is a critical need for alternative methods that enable correlation of gene expression profiles with the phenotypic characteristics of adherent single cells.

In this study, we designed a photoresponsive cell culture surface composed of two materials to enable the construction of single-cell arrays, time-course morphological tracing, and selective release of individual cells for downstream gene expression analysis (Figure 1). We have recently developed a photoactivatable cell-anchoring material for single-cell trapping at high density; this material, lipid-conjugated poly(ethylene glycol) (PEG) **1**, is converted from the double-lipid-conjugated form to the single-lipid form (**1a**) via the photocleavage of one lipid moiety on the substrate surface (Figure 1A) [26]. This conversion enabled selective cell trapping in light-irradiated regions through interactions between the cellular lipid bilayer membranes and the single lipid moiety exposed on the substrate surface (Figure 1A). After cell trapping, adherent cells can freely adhere to and spread on the underlying surface without being spatially restricted by the cell-anchoring material, thereby allowing them to exhibit their cellular functions, such as migration and proliferation [26]. For selective cell release after microscopic monitoring of adherent cells, a photodegradable hydrogel layer was used as the basal cell-adhesive surface under the cell-trapping material (Figure 1B).

**Figure 1.**
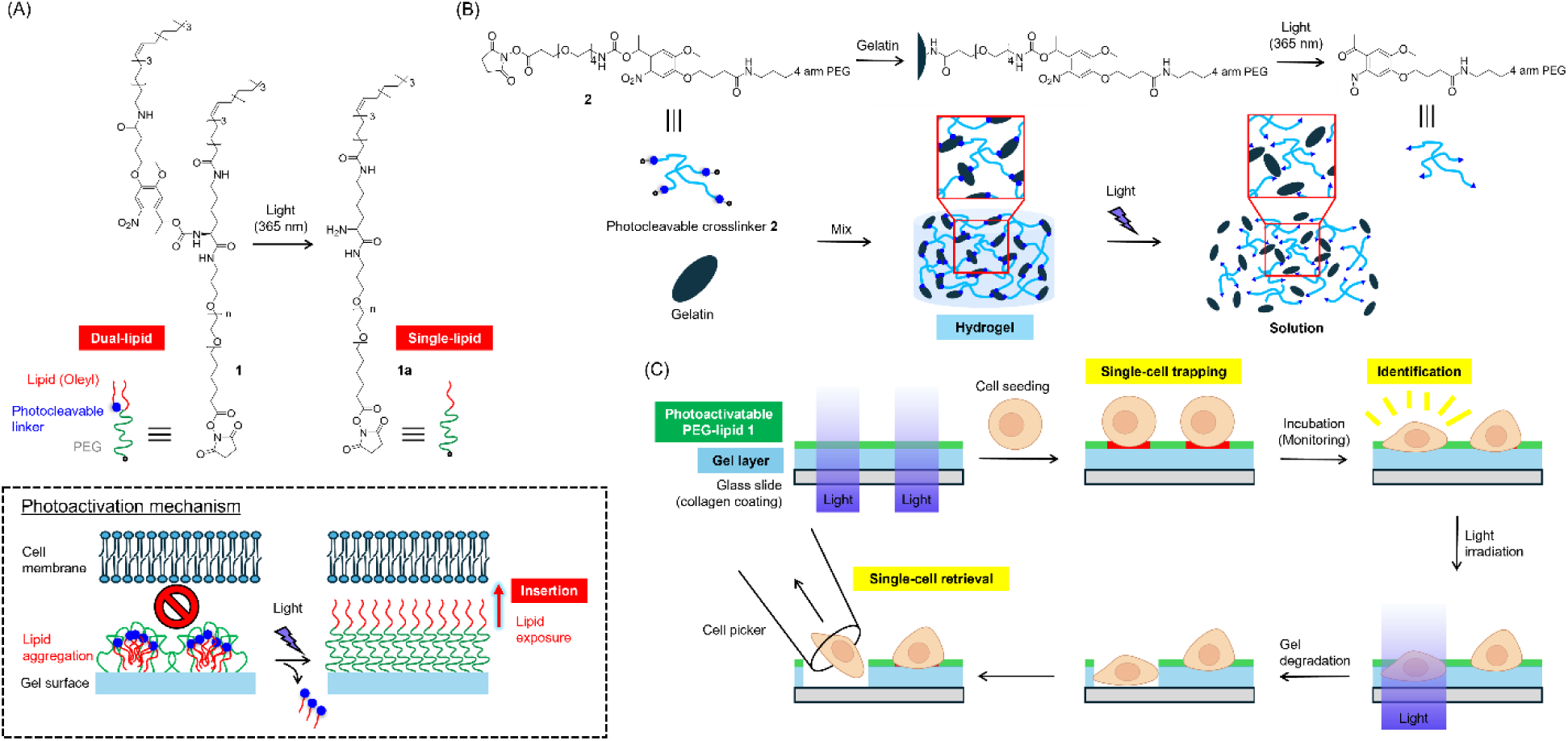
Schematic illustration of dual photoresponsive hydrogel surfaces for single-cell analysis. (A) Structural changes of the photoactivatable cell-trapping material **1** before and after light irradiation (365 nm) and illustration of the photoactivation mechanism. (B) Chemical reactions and schematic illustrations showing the formation and degradation of the photo-degradable hydrogel composed of gelatin and the synthesized photocleavable crosslinker **2**. (C) Schematic illustration of the photoresponsive gel-layer surface for cell trapping, image-based analysis, and cell release. The initial light irradiation activates the photoactivatable cell-anchoring material **1**, enabling single-cell-level trapping. After incubation and monitoring, the photodegradable gel layer beneath the target cells is irradiated to induce local degradation, allowing the cells to be released and then collected for downstream genetic analysis.

Hydrogels are highly attractive materials for artificial cell culture platforms because they can mimic the natural extracellular matrix (ECM) [27]. Furthermore, photodegradable hydrogels have been reported to enable selective release of adherent cells through light-induced degradation without causing cellular damage [28–30]. Over the past two decades, various platform technologies using photodegradable hydrogels have been developed for cell patterning and release [31–34]. However, to the best of our knowledge, no method has been reported for constructing precisely defined single-cell arrays on photodegradable hydrogels in a light-guided manner. In this study, a photodegradable hydrogel layer was prepared by crosslinking gelatin with photocleavable four-branched polymeric crosslinker **2**. As shown in Figure 1C, by modifying the surface of the photodegradable gel layer with PEG-lipid **1**, cells were trapped at high density with uniform spacing by irradiating a designed array of light spots. Consequently, in contrast to random cell seeding, the individual cells were not excessively close to each other on the hydrogel surface, even at a high density, thereby preventing cell–cell contact during monitoring and allowing each cell to be retrieved individually without collecting adjacent cells. The trapped adherent cells were monitored using time-lapse microscopy as they adhered to the gel layer surface, changing their morphology. Through high-throughput image analysis, cells exhibiting a specific phenotype were rapidly identified and noninvasively released from the surface via gel degradation induced by secondary selective light irradiation (Figure 1C). The released single cells were collected individually using a microcapillary and then were subjected to genetic analysis, including single-cell real-time quantitative PCR and single-cell RNA sequencing. Using this workflow, we demonstrated the ability to identify heterogeneity among invasive model cancer cells using time-lapse microscopy and to examine the changes in the gene expression profiles associated with their morphological characteristics.

## Results

### Synthesis and Characterization of Photodegradable Gel Surfaces

The photodegradable hydrogel layer was fabricated by mixing gelatin with photocleavable polymeric crosslinker **2**, consisting of four-arm PEG functionalized with amine-reactive N-hydroxysuccinimide (NHS) esters at each arm terminus via photocleavable 6-nitroveratryloxycarbonyl (Nvoc) linkers (Figure 1B). NHS esters react with the amine groups of the gelatin molecules to form crosslinks, resulting in gelation. Upon exposure to light, the Nvoc linkers are cleaved, leading to hydrogel degradation (Figure 1B). Photocleavable crosslinker **2** was synthesized in five steps (Figure S1), and the representative peaks in the ^1^H NMR spectra were assigned (see Supporting Information). A premixed solution of gelatin and photocleavable crosslinker **2** was extended onto a thin layer on a glass slide and allowed to form a thin gel layer.

We examined whether cell-trapping materials could be immobilized on the gel surface by utilizing the remaining amine groups of gelatins after cross-linking. Non-photoresponsive PEG-lipid **3** was used as a model cell-trapping material (Figure 2A). In this experiment, the entire surface of the green fluorescence-labeled gel layer was modified with **3**. PEG-lipids can rapidly anchor to a wide range of cell types (within 5 min) through interactions with the lipid bilayer membranes of cell surfaces regardless of their cellular adhesiveness [35, 36]. Accordingly, after cell seeding onto the gel layer surface, cells anchored via interactions with PEG-lipids can be retained, whereas non-anchored cells are washed away by rinsing the substrate surface within 10 min, as stable biological cellular adhesion generally requires a longer time [37–39]. In this study, a microchannel system was employed to ensure uniform rinsing of the surface, thereby removing nonspecific and light-induced released cells, as described in our previous work [15, 40]. A cover slide with microchannels was placed on the gel layer-coated glass slide and secured with a fixture (Figure S2). HeLa cells (human cervical carcinoma cell line) were introduced from the microchannel inlet and incubated for 10 min. After rinsing, the cells were confirmed to have anchored to the gel layer (Figure 2C). This result strongly indicates that a simple treatment of the gel layer with PEG-lipids enables surface modification and subsequent cell anchoring to the gel layer surfaces.

**Figure 2.**
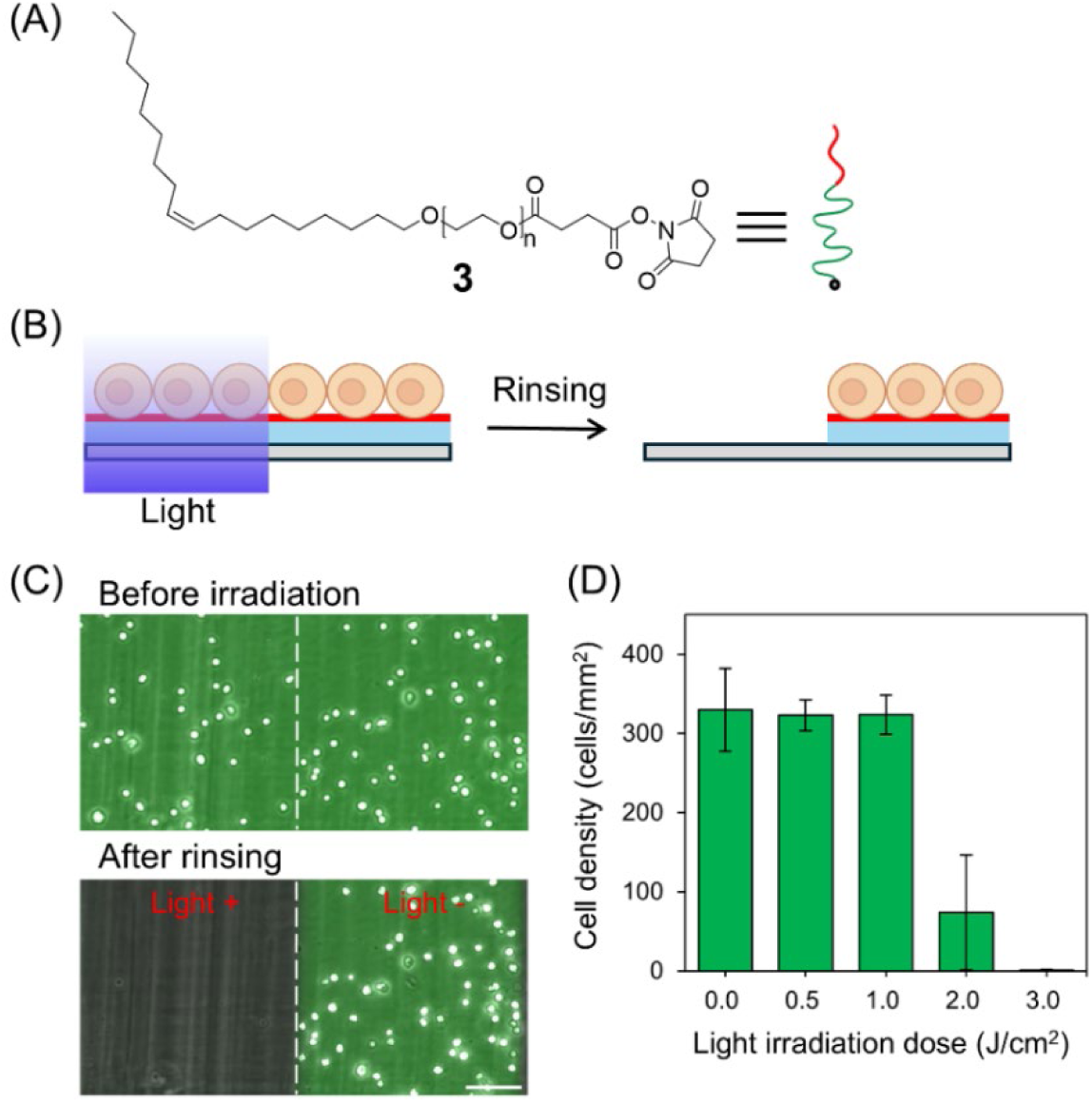
Light-induced degradation of cell-anchored gel layers. (A) Chemical structure of poly(ethylene glycol)–lipid **3** used for cell trapping. (B) Schematic illustration of light-induced gel degradation and cell release. (C) Fluorescence microscopic images of anchored HeLa cells on fluorescein-stained gel layers (green) before and after light irradiation-induced gel degradation followed by rinsing. Scale bar: 100 μm. (D) Cell densities after exposure to varying doses of light. Values and error bars represent the mean ± standard error (n = 9).

To examine the light-induced release of the anchored cells via gel photodegradation, portions of the gel layer were exposed to ultraviolet light (365 nm) at varying light doses (Figure 2B). After light exposure and subsequent rinsing, both the gel and anchored cells selectively disappeared from the light-irradiated regions (Figure 2C and S3). This result clearly demonstrates that the anchored cells can be selectively released through light-induced gel degradation. Furthermore, the cell density on the gel layer increased at a dose of 2 J/cm^2^ and decreased to nearly zero at a dose of 3 J/cm^2^ (Figure 2D). These light doses (≤ 5 J/cm^2^) have not been found to affect cell viability in previous studies on photoresponsive cell manipulation materials [40, 41]. Thus, the present PEG-lipid-modified gel surface enables cell trapping and release under non-cytotoxic conditions.

### Light-Induced Cell Patterning and Release on Dual Photoresponsive Gel Surfaces

To achieve both spatially controlled cell trapping and selective cell release, we modified the photodegradable gel layer with photoactivatable PEG-lipid **1** (Figure 1A). In this design, photoactivatable PEG lipids were selectively activated in the light-exposed region upon localized light exposure. Subsequently, the seeded cells were rapidly anchored through interactions with the activated PEG-lipid, resulting in the formation of well-defined cell patterns after washing away the non-anchored cells. Furthermore, the patterned cells were released by a second light exposure, followed by rinsing (Figure 3A). However, as described above, the photodegradable gel layer was partially dissolved upon exposure to a light dose of 2 J/cm^2^. Accordingly, the initial light exposure for PEG-lipid activation should be performed at doses below 2 J/cm^2^ to anchor the cells without inducing gel degradation. When the gel layer was exposed to line-patterned light at a dose of 0.5–1 J/cm^2^, the density of anchored cells in the irradiated regions increased proportionally with the light dose and reached nearly full confluence at 1 J/cm^2^ (Figure S4). This behavior can be attributed to the fact that cell anchoring mediated by the photoactivatable PEG-lipid is more photosensitive than the photodegradation of the hydrogel, even though both molecular systems utilize the same photocleavable linker. These results provided strong evidence that the present approach enabled the development of a gel layer capable of both cell trapping and release. Indeed, on the gel layer surface, HeLa cells were selectively anchored to light-irradiated regions at a light dose of 1 J/cm^2^ (Figure 3B, left) and subsequently released via gel photodegradation induced by the second dose of light irradiation at 5 J/cm^2^ followed by rinsing (Figure 3B, right). This second irradiation step led to the selective release of the anchored cells from the irradiated regions, whereas the cells in the non-irradiated regions remained unaffected. Thus, the dual-photoresponsive gel layer achieved precise spatial control of both cell trapping and release.

**Figure 3.**
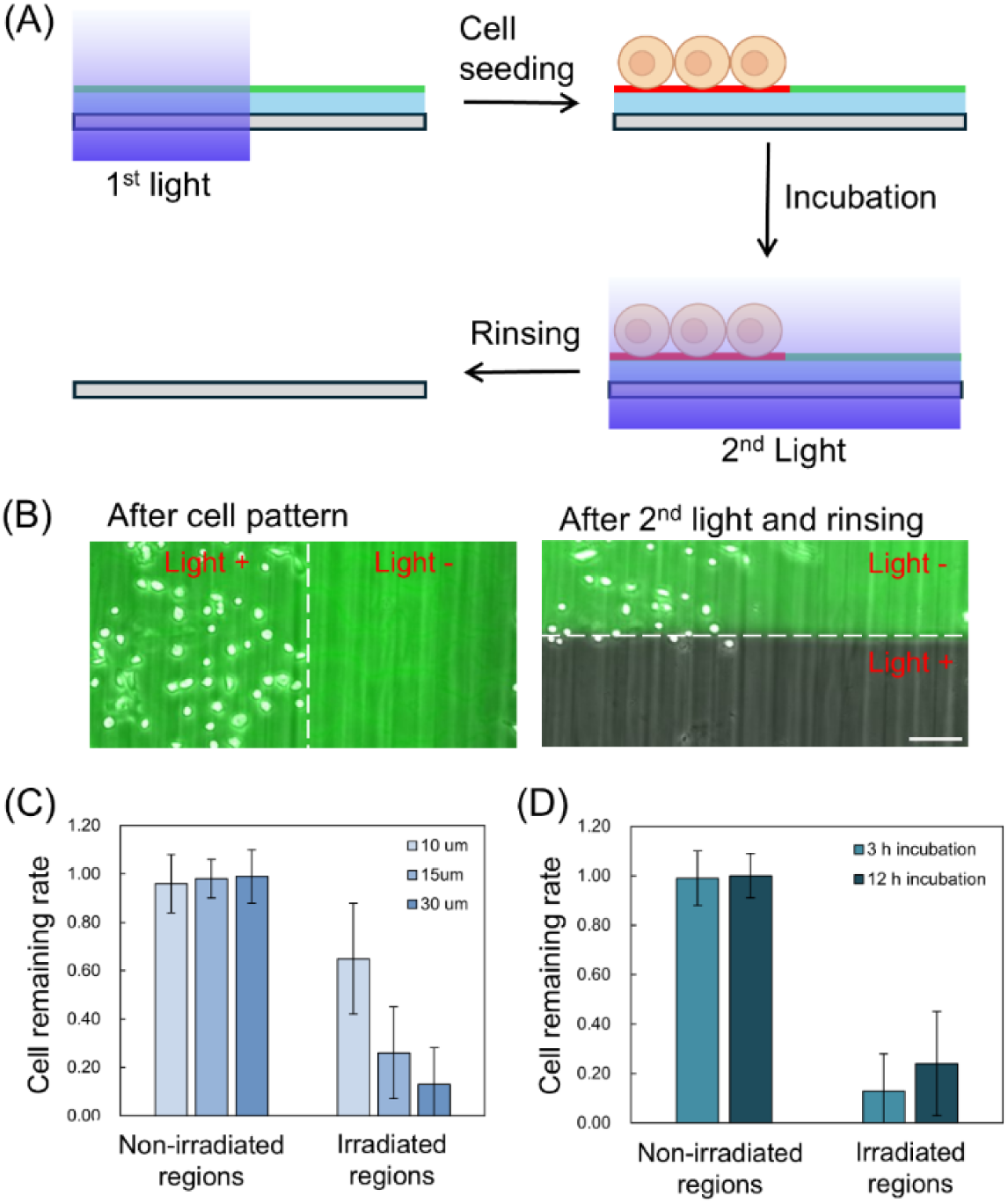
Light-induced cell trapping and cell release. (A) Schematic illustration of light-induced cell trapping and release. (B) Fluorescence microscopy images of anchored HeLa cells on fluorescein-stained gel layers (green) in regions with or without the first and second light-irradiation treatments. *Left*: after cell patterning; *Right*: after the second light-irradiation treatment followed by rinsing. Scale bar: 100 μm. (C) Cell-remaining rates after second round of light irradiation in non-irradiated and irradiated regions on gel layers with different thicknesses. Values and error bars represent the mean ± standard error (n = 36). (D) Cell-remaining rates after second round of light irradiation in non-irradiated and irradiated regions after incubation for 3 h and 12 h. Values and error bars represent the mean ± standard error (n = 36).

We further investigated how the culture period between cell trapping and release affected the cell release efficiency. The light-anchored HeLa cells were cultured on the gel layer for 3 h and subsequently released by light irradiation. Surprisingly, for the 10-µm-thick gel layer used in the experiments described above, most cells remained on the surface even after light irradiation followed by rinsing (Figure 3C). Based on this observation, we hypothesized that cells could interact with the underlying substrate surface rather than with the gel layer, resulting in low cell release efficiency. To prevent such interactions between the cells and substrate surface, the thickness of the gel layer was increased, and it was found that the rate of cell retention decreased with increasing gel thickness (Figure 3C). On 30-µm-thick gel layers, the cell release efficiency after 3 h of culture reached approximately 85%. In addition, we evaluated the effect of incubation duration by extending the time period of cell culture on the gel layer from 3 h to 12 h. Extending the culture period led to an increase in the rate of cell retention after light irradiation; however, approximately 75% of the cells could still be successfully released from the irradiated regions (Figure 3D). These results suggest that the present method for cell trapping and release is applicable not only for short-term, but also for long-term cell cultures. Long-term time-lapse observation of dynamic cell behavior is a powerful approach for identification of unique cells that are difficult to distinguish within a cell population prior to cell retrieval, thereby expanding the range of applicable cellular events.

### Light-Induced Single-Cell Isolation from Dual Photoresponsive Gel Surfaces

We demonstrated that the gel layer surface can be used for image-based single-cell isolation. The surface was exposed to an array of light spots with a single cell diameter, as reported in our previous studies [15, 26, 39]. Individual HeLa cells were anchored at each light-irradiated spot with high efficiency at the optimal spot size (single-cell spot rate, 84%) (Figure 4). Furthermore, biological adhesion of the anchored cells on the surface was examined by fluorescent staining of both actin filaments and focal adhesions after culture for 13 h. In individual anchored cells, well-developed actin filaments and focal adhesion were observed near the plasma membrane along with filopodial protrusions at the cell periphery (Figure S5). These observations strongly indicated that the anchored cells biologically adhered to the gel layer. Comparison with the control photodegradable gel layer without PEG-lipid modification revealed no marked differences in the adhesion state (Figure S5). Accordingly, PEG-lipid modification did not significantly influence cellular adhesion under present conditions, which is consistent with our previous reports on photoresponsive PEG-lipid-modified surfaces [26, 39]. Thus, the present gel layer surface enabled construction of a fine single-cell array of adherent cells in which individual cells adhered to the surface and exhibited normal cellular behavior.

**Figure 4.**
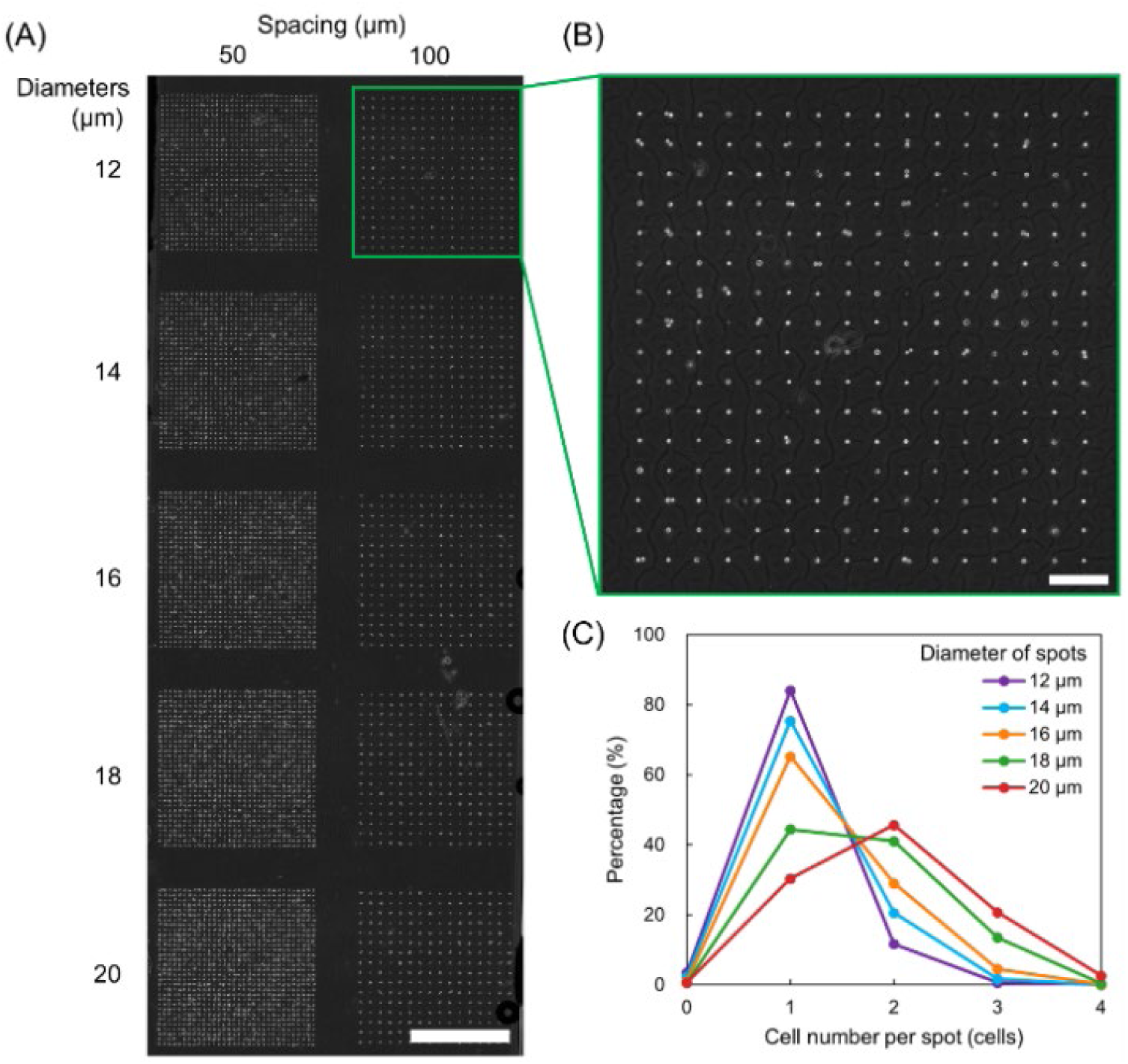
Single-cell array formation on dual photoresponsive gel layers. (A) Bright-field microscopy images of arrayed HeLa cells on the gel layers after exposure to arrays of light spots with varying diameters (12 – 20 μm) and two different inter-spot spacings (50 and 100 μm). Scale bar: 1000 μm. (B) Enlarged images of arrayed HeLa cells (spot diameter: 12 μm; inter-spot spacing: 100 μm). Scale bar: 200 μm. (C) Percentages of spots capturing different numbers of cells per spot in the cell array with an inter-spot spacing of 100 μm. The percentages for spots with varying diameters are plotted.

To clearly demonstrate image-based isolation of target cells, we prepared a single-cell array composed of a mixture of HeLa cells and enhanced green fluorescent protein (EGFP)-expressing HeLa cells (HeLa–EGFP) on a red fluorescence-labeled gel layer surface. Individual cells were identified as either HeLa or HeLa–EGFP based on their green fluorescence signals in the microscopic images. First, a randomly selected HeLa cell was exposed to a localized spot of light to induce detachment from the surface and then retrieved using a capillary-based cell picking system (Figure 5A), followed by dispensing into a microtube. This operation cycle was repeated sequentially, and HeLa–EGFP cells were retrieved from the surface. After picking, the cells at the light-irradiated spot disappeared and the fluorescence of the gel layer vanished (Figure 5B). These results strongly indicated that each light-irradiated cell could be detached via the photodegradation of the gel layer and subsequently retrieved by aspiration using a capillary.

**Figure 5.**
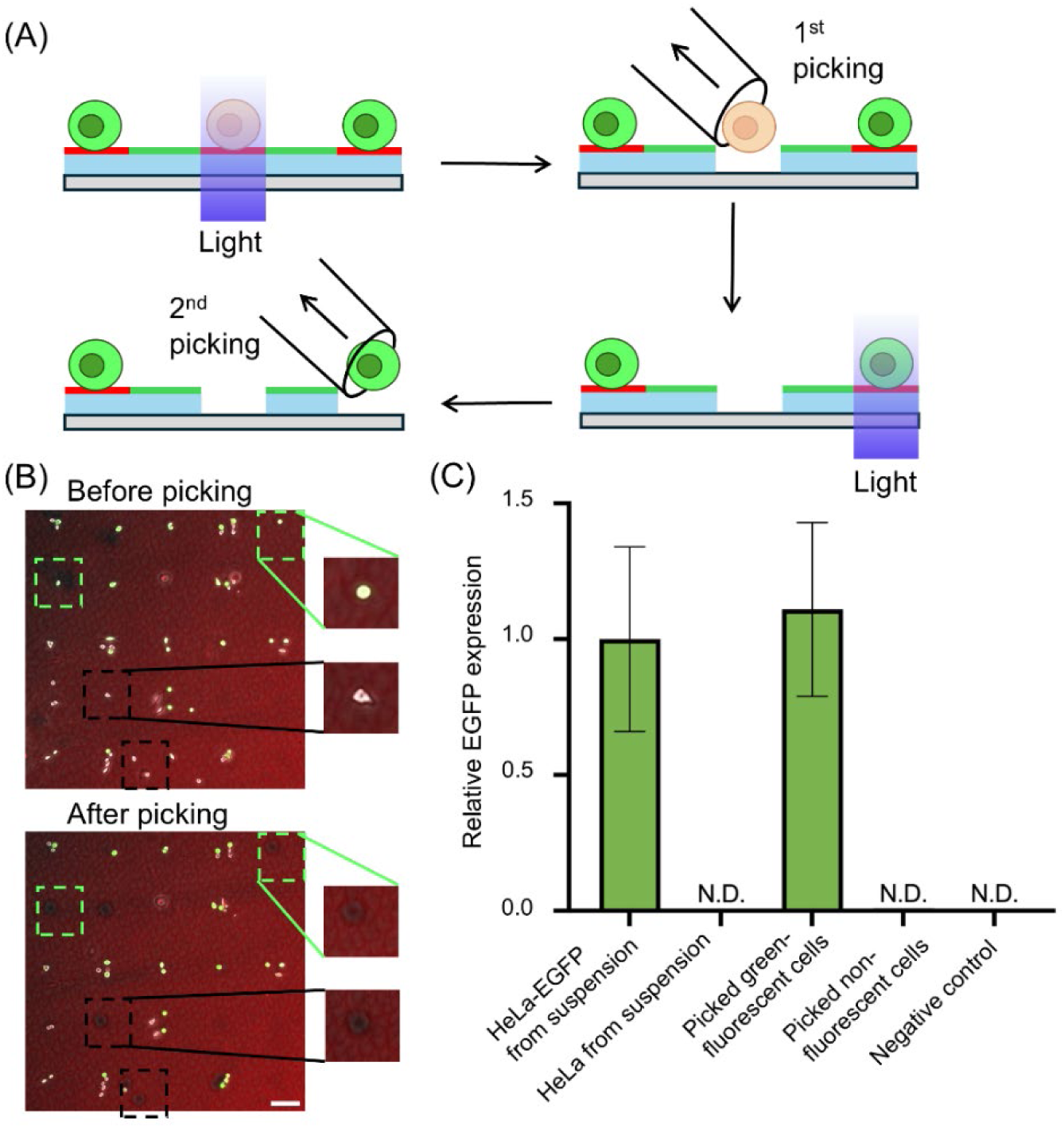
Light-induced single-cell retrieval and gene expression analysis. (A) Schematic illustration of light-induced single-cell release and collection. (B) Fluorescence microscopy image of anchored HeLa and HeLa-EGFP cells on a rhodamine-stained gel layer (Red) before light-induced cell release and after cell picking. Scale bar: 100 μm. (C) Relative EGFP expression levels normalized to GAPDH expression. HeLa or HeLa-EGFP from suspension: the single cells collected from the diluted cell suspensions using a cell picker; Picked cells: green-fluorescent or non-fluorescent single cells collected from the single-cell array by cell picking; Negative control: PBS. Values and error bars represent the mean ± standard error (n = 3).

The retrieved single cells were characterized based on specific gene expression profiles using single-cell reverse transcription quantitative PCR (RT-qPCR) analysis. Prior to the single-cell analysis, we confirmed that the cells retrieved from the gel layer surface could be analyzed by RT-qPCR using a population of anchored cells. HeLa and HeLa-EGFP cells were independently anchored in linear patterns across the entire surface of the gel layer. Portions of the patterned surfaces with varying areas were then exposed to light to induce cell release, followed by the retrieval of the released cells by pipetting. RT-qPCR analysis of the retrieved cell populations revealed that EGFP transcripts were selectively detected only in cells retrieved from the HeLa-EGFP patterns (Figure S6). These results demonstrate that the present RT-qPCR protocol is suitable for quantitative gene expression analysis of cells retrieved from the gel layer surface. Based on this validation, the retrieved single cells were analyzed using the same procedure. The EGFP transcripts were selectively detected in the retrieved green-fluorescent cells (Figure 5C). In addition, the relative transcriptional level of EGFP normalized to that of an endogenous control gene was comparable to that observed in intact HeLa-EGFP cells. These results clearly indicated that individual target cells could be accurately collected from the gel layer surface in accordance with the phenotypes identified by microscopy observation without causing significant transcriptional perturbations.

### Integrated Single-Cell Analysis of Cellular Morphology and Gene Expression

The gel layer surface was applied to perform integrated single-cell analysis of cellular morphology and gene expression. The gel layer consists of cross-linked gelatin, and gelatin-based hydrogels are well-known to provide an ECM-like cell culture environment [42, 43]. Furthermore, cells sense and respond to the mechanical properties of the ECM environment, and these mechanotransduction pathways critically influence cell morphology, adhesion, and gene expression [44, 45]. Therefore, HT1080 cells (a human fibrosarcoma cell line) were used as the target for the integrated analysis in this experiment because of their high motility and metastatic potential mediated by interactions with the ECM [46]. Individual HT1080 cells were retrieved from a single-cell array based on their morphology (Figure 6A). After approximately 14 h of incubation, a subset of HT1080 cells exhibited a spread morphology, whereas others retained a round shape (Figure 6B). Based on these microscopic images, the cells were visually classified into two groups, released from the gel layer surface by light irradiation, and individually collected into microtubes using a cell picker. The retrieved cells were analyzed by RT-qPCR to quantify the expression of several genes involved in cell adhesion and invasion. For matrix metalloproteinase-9 (MMP9), an enzyme associated with ECM degradation and cancer cell invasiveness, significantly higher MMP9 transcript levels were observed in the cells exhibiting spread morphology than in the cells with round morphology (Figure 6C).

**Figure 6.**
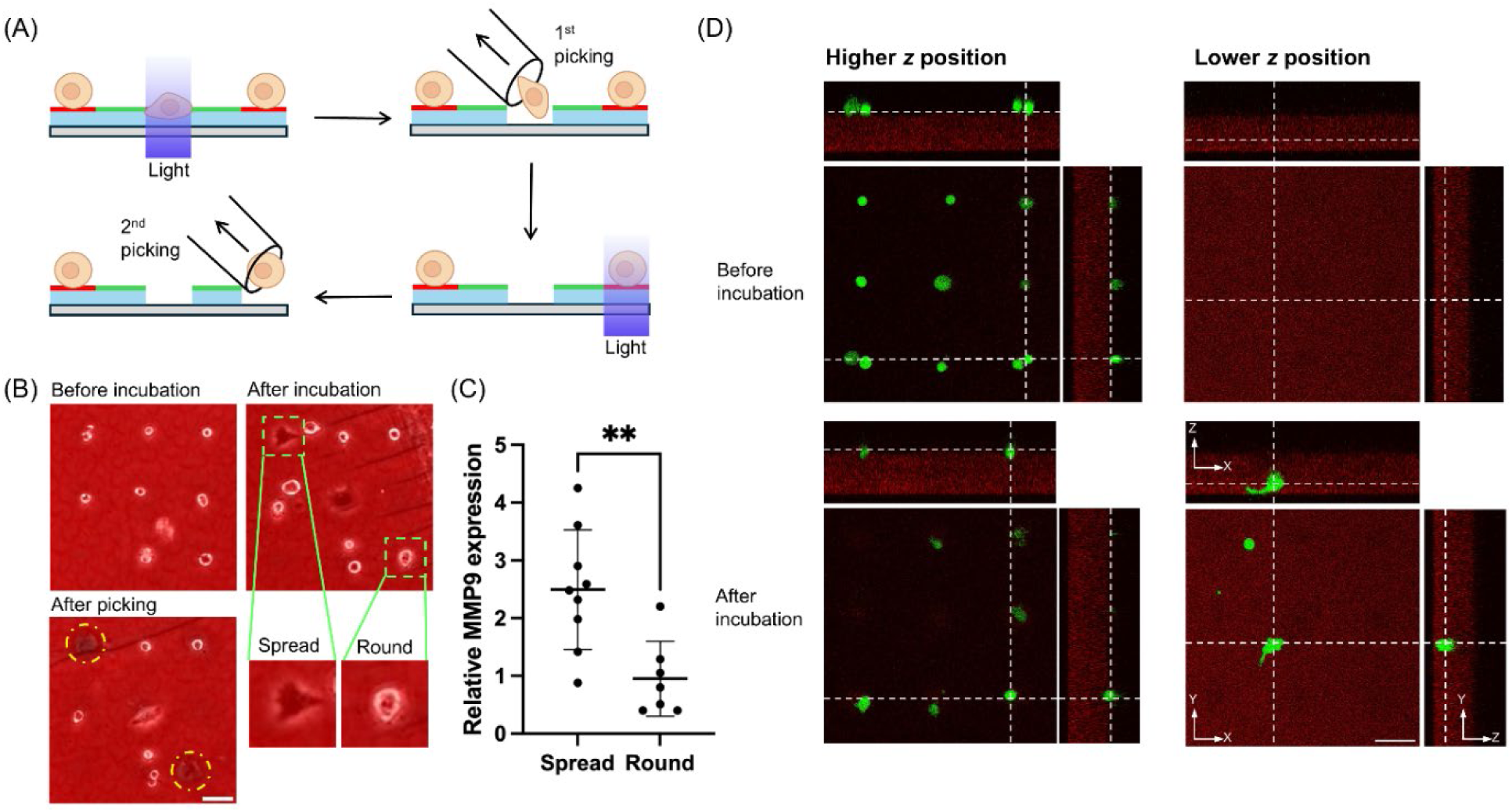
Morphology-based single-cell retrieval and gene expression analysis. (A) Schematic illustration of morphology-based single-cell release and collection. (B) Fluorescence microscopy images of anchored HT1080 cells on a rhodamine-stained gel layer (Red) before incubation, after incubation, and after single-cell picking using a cell picker. Scale bar: 100 μm. (C) Relative MMP9 expression levels normalized to GAPDH expression. Spread: cells exhibiting a spread morphology; Round: cells exhibiting a round morphology. (D) Confocal fluorescence microscopy images of anchored HT1080 cells in the *x-y*, *x-z* and *y-z* planes. Images were acquired at higher and lower *z* positions before and after incubation. HT1080 cells were stained by CellTracker Green CMFDA (Green). Scale bar: 100 μm

In addition, using confocal microscopy, we monitored the changes in the z-position of the fluorescently labeled HT1080 cells before and after incubation. As shown in Figure 6D, all cells were located on the surface of the gel layer before incubation. After incubation, some cells invaded the gel layer, whereas others remained on the gel surface. Based on these observations, we hypothesized that cells with higher MMP9 expression are more capable of degrading the surrounding gelatin-based gel, thereby creating sufficient space for spreading. By contrast, cells with lower MMP9 expression may be unable to adequately remodel their microenvironment, and therefore remain rounded. These results demonstrate that morphology-based single-cell identification can reveal heterogeneity in gene expression, highlighting the potential of phenotype-guided integration analysis for understanding cellular behavior at the molecular level.

Finally, to demonstrate the potential of the present system, we performed integrated analysis that combined automated time-course morphological profiling and gene expression analysis. Subtle morphological differences between individual cells are difficult to resolve via visual inspection and time-lapse imaging generates a large number of single-cell images, rendering manual quantification impractical. Therefore, we implemented a high-throughput, real-time image-processing pipeline that enabled automated recognition, tracking, and quantitative measurement of individual cells in single-cell arrays imaged every 5 min over a 14-h incubation period (Figure. 7A). Cells cultured on a single-cell array were characterized using time-course morphological fingerprints defined by statistical descriptors derived from multiple morphological features established in previous morphology-based cell evaluation studies [12, 47]. Specifically, 26 descriptors consisting of the means and standard deviations of 13 basal morphological features over the 14-h observation period were used to quantify the temporal morphological dynamics. This approach enabled objective discrimination between the cells exhibiting active morphological changes and those displaying relatively stable morphologies. Representative examples of these time-course morphological fingerprints are shown in Figure 7B.

**Figure 7.**
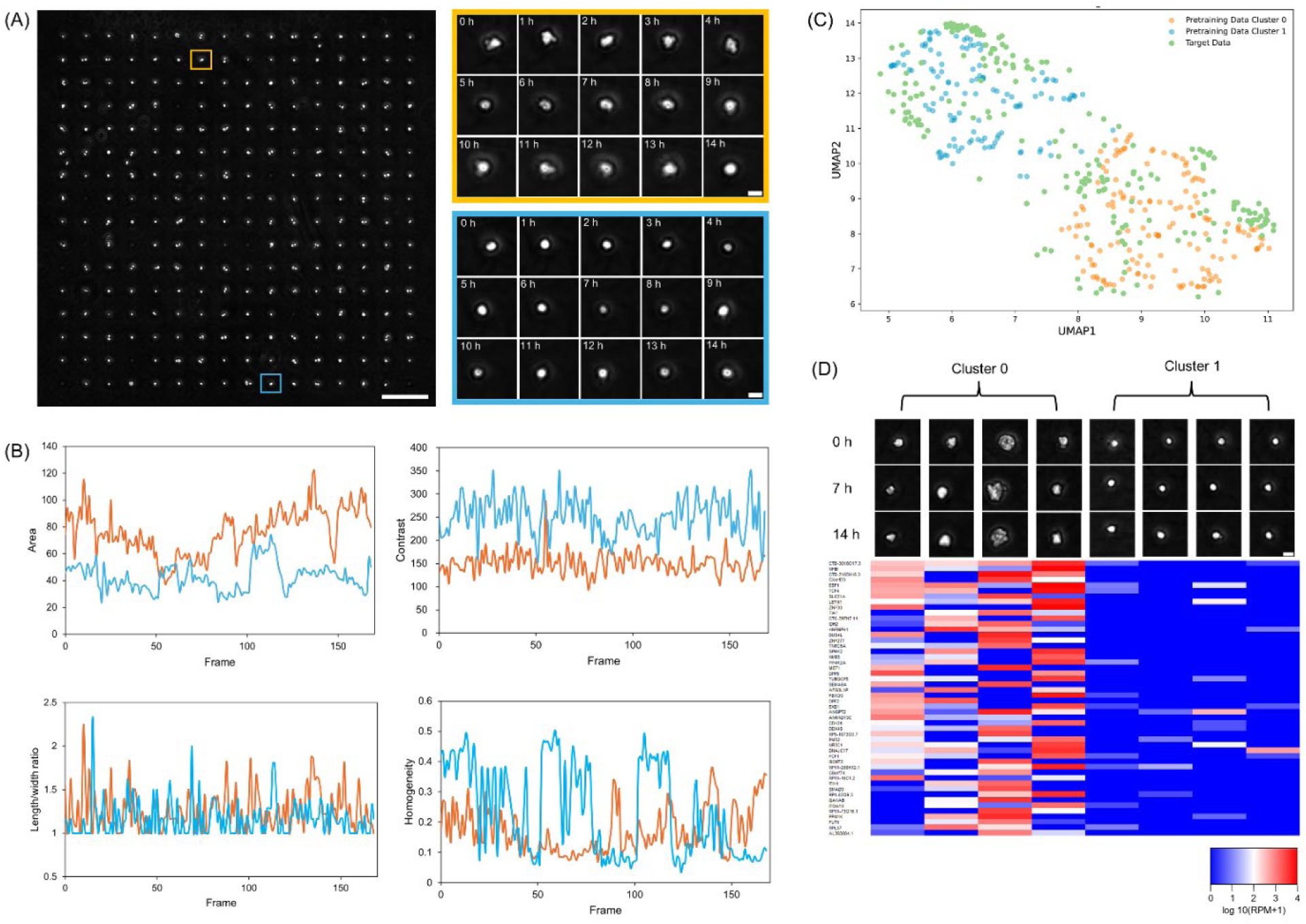
Integrated analysis of single-cell morphological profiling and gene expression profiling. (A) Microscopy images of a single-cell array of HT1080 cells on the gel layer surface. Representative cells exhibiting active or subtle morphological changes are high-lighted by pale orange and pale blue squares, respectively. Enlarged time-lapse images of these two representative cells are shown. Scale bar: 200 μm (*left*); 20 μm (*right*). (B) Plots showing the time evolution of the area, contrast, length-to-width ratio, and homogeneity for the two representative cells shown in A. Pale orange: cells exhibiting active morphological changes; pale blue: cells exhibiting subtle morphological changes. (C) UMAP visualization of single-cell morphological profiles of pretrained reference cells and target cells. Pretrained cells are classified into two major clusters; cluster 0, active morphological change (pale orange); cluster 1, subtle morphological change (pale blue). Target cell data (pale green) are overlaid onto the reference UMAP space. (D) Time-lapse microscopy images of representative cells from each cluster and their corresponding transcriptomic analysis results. Scale bar: 20 μm.

Using these multidimensional temporal descriptors, the HT1080 cells cultured on the array were classified into two major morphological clusters by UMAP (Uniform Manifold Approximation and Projection) analysis (Figure 7C). Using these pretrained reference cells, newly cultured cells on the array were categorized in real time as target cells according to their cluster assignment. Importantly, this real-time quantitative morphological profiling enabled not only the systematic classification of all cells on the array according to their morphological change potential, but also the quantitative identification and physical retrieval of the target cells directly from the single-cell array. This framework facilitated the visualization and analysis of heterogeneous single-cell subpopulations in a single-cell array.

To assess whether these morphological differences reflected the underlying transcriptional heterogeneity, we selectively retrieved individual cells from each cluster using light-induced cell release for single-cell transcriptome analysis. Based on the UMAP generated from the target dataset, representative single cells from each cluster were retrieved and subjected to transcriptome analysis by single-cell RNA sequencing (scRNA-seq). As shown in Figure 7D, when mapped back onto their corresponding morphological clusters, the resulting gene expression data revealed both inter- and intra-cluster heterogeneity at the transcriptomic level. Despite this heterogeneity, we identified genes that were commonly upregulated in cells within cluster 0 and were characterized by relatively active morphological changes. For instance, transcription factor 4 (TCF4) is upregulated in three of the four cells in cluster 0 and has been reported to be overexpressed in highly invasive cancer cells [48]. Moreover, TCF4 is an important regulator of epithelial–mesenchymal transition (EMT). Thus, the upregulation of previously reported genes associated with morphological changes supports the validity of the integrated single-cell analysis. Concurrently, we detected the upregulation of genes whose roles in morphological changes have not yet been characterized. One such RNA (lncRNA), CTD-3018O17.3, was upregulated in all cells within cluster 0. Although the function of this gene is not fully understood, it may represent a novel candidate for regulation of cancer invasion. These results underscore the potential of our platform for bridging the gap between imaging-based phenotypic characterization and downstream gene expression profiling. Importantly, the ability to distinguish cells based on subtle predefined behaviors prior to transcriptomic profiling provides a powerful strategy for analysis of dynamic cellular states that would be averaged using conventional bulk or random-single-cell-sampling approaches.

## Discussion

In this study, we developed a dual photoresponsive gel-layer surface that enables both single-cell patterning and release through controlled photoirradiation. Compared with conventional surfaces for single-cell manipulation, this platform offers several notable advantages. First, both the capture and release of single cells can be achieved by two times’ light irradiation. Unlike previously reported photodegradable gel surfaces, on which cells can only be randomly seeded, single cells can be arrayed at user-defined fine intervals on the present gel-layer surface using photoactivatable PEG-lipids. Therefore, depending on the target cells to be observed and their events of interest, cells can be arranged at the maximum achievable density by defining a minimal spacing that prevents cell overlaps during both observation and cell retrieval. In this study, the spacing was set to 100 μm because long-term time-lapse migration of HT1080 cells was one of the target events to be observed (Figure 6). As a result, high-throughput single-cell observation was achieved on photodegradable gel surfaces for the first time. Using the resulting large dataset, cells could be automatically clustered by UMAP, enabling discrimination of even subtle differences in morphological characteristics of cells from the same cell line (Figure 6).

In addition, unlike most conventional single-cell arrays that confine cells within narrow microstructures, cells trapped on the present gel-layer surface were able to freely spread and migrate during time-lapse observation without physical constraints. Furthermore, this gelatin-based gel layer supports cell adhesion. In conventional *in vitro* culture, cells are typically maintained on rigid plastic or glass substrates, which provide a mechanical environment that is orders of magnitude stiffer than native tissues. In contrast, ECM *in vivo* exhibits complex mechanical behaviors, including viscoelasticity and nonlinear elasticity, which more faithfully reflect the conditions experienced by cells within tissues [49, 50]. Accordingly, in some cases, cells cultured on ECM-mimicking soft matrices display distinct morphologies, cytoskeletal organization, and mechanosensitive signaling compared with cells on rigid two-dimensional substrates [49, 51]. Notably, some HT1080 cells were observed to invade into the gel layer, and the heterogeneity in invasion ability was detected based on morphological features (Figure 5). Cell invasion has traditionally been evaluated using Boyden chamber–based assays, which primarily assess invasion at an endpoint [52]. In contrast, the present gel layer–based platform enables direct, time-lapse visualization of invasion behavior at the single-cell level. Together, these results suggest that the present single-cell array based on a soft ECM-like hydrogel can more effectively recapitulate certain *in vivo* cellular behaviors that are difficult to achieve using conventional rigid substrates.

After time-lapse observation, individual adherent cells on the gel-layer surface could be selectively collected through light-induced detachment for downstream gene expression analysis, enabling a direct link between dynamic morphological features and gene expression profiles at the single-cell level. As with reported photodegradable gel surfaces [28, 29], cells were retrieved alive via local gel degradation. Gene expression analysis by RT-qPCR confirmed that light-induced cell retrieval is compatible with single-cell RNA quantification without causing significant transcriptional perturbation, likely due to minimal cell damage (Figure 4). Accordingly, a distinct cellular morphological feature could be linked to the expression of an associated gene. In this study, a spread morphology on the gel layer was found to correlate with MMP9 expression. Based on this finding, cellular invasion into the gel layer was further examined by confocal microscopy, leading to the inference that MMP9 is involved in the mechanism underlying the observed morphological features. Moreover, subtle dynamic morphological features distinguished by image analysis of a large dataset were also linked to gene expression profiles (Figure 6D). In this trial, cells exhibiting active morphological changes showed upregulation of genes previously reported to be associated with morphology, as well as genes whose roles in morphological changes remain unclear. It is expected that finer clustering in the morphological profiling will clarify cluster-specific upregulation and downregulation in gene expression. Nevertheless, this study provides proof of principle that the present photoresponsive gel-layer surface is a promising platform for integrated single-cell analysis of cellular behavior and gene expression.

## Conclusions

We have developed a dual-photoresponsive cell culture surface that enables high-density single-cell patterning, time-lapse morphological analysis, and selective cell retrieval for downstream gene expression profiling. By integrating a photoactivatable PEG-lipid with a photodegradable hydrogel layer, the surface first allows precise spatial trapping and release of individual cells via light irradiation. This surface supports cell spreading and migration in an ECM-like environment, enabling the capture of dynamic and heterogeneous cellular behaviors. Coupled with high-throughput imaging and gene analysis, it establishes a direct link between single-cell morphology and transcriptional alterations. This approach provides a versatile framework for integrated single-cell analysis and offers potential for uncovering phenotype–gene expression relationships in complex biological systems. Furthermore, this dual photoresponsive surface is applicable not only to integrated analysis with single-cell transcriptomics but also to other single-cell omics analysis, such as proteomics [53]. In particular, compared with conventional enzymatic harvesting of adherent cells, the present light-induced cell retrieval offers significant advantages in that adherent cells can be collected without enzymatic digestion of cell-surface proteins, which is especially beneficial for single-cell proteomics. In addition, by varying the size and pattern of the irradiated light, both single-cell density and the number of cells per spot can be precisely controlled. By repeatedly applying light irradiation and cell anchoring, different cell types can be arranged in defined configurations on the present photodegradable surfaces, enabling unique single-cell analysis of cell-cell communication [15, 26], followed by gene expression analysis of individual cells to identify key molecules in the interactions of interest. Such flexibility has the potential to facilitate new types of experiments in drug screening and disease modeling by enabling the analysis of dynamic cellular behaviors and their molecular signatures. Beyond cancer cells, this approach may also be applicable to other cell types, such as neurons or immune cells, thereby offering broader utility in basic and translational research. Accordingly, the present dual photoresponsive gel-layer surface provides a versatile platform for single-cell studies across bioscience, bioengineering, and medical applications.

## Materials and methods

### Synthesis of a Photocleavable Crosslinker

Synthetic protocols are described in detail in the Supporting Information. Briefly, the photocleavable crosslinker **2** was synthesized by functionalizing the termini of a four-arm PEG (Mw: 10000; Sunbright PTE-100PA, NOF Co., Tokyo, Japan) with a photocleavable Nvoc linker. Carboxyl groups were then introduced at each functionalized terminus and converted to NHS esters for crosslinking with the amine groups of gelatin during gel formation.

### Cell Culture

Hela cells were cultured in low-glucose DMEM (Wako) supplemented with 10 % bovine serum (FBS; Gibco), 100 u/mL penicillin, and 100 μg/mL streptomycin. Hela-EGFP cells were cultured in high-glucose DMEM (Wako) supplemented with 10 % FBS, 0.1 mM MEM non-essential amino acids (NEAA), 100 U/mL penicillin, and 100 μg/mL streptomycin. HT1080 cells were cultured in low-glucose DMEM supplemented with 10 % FBS, 100 u/mL penicillin, and 100 μg/mL streptomycin. All cells were maintained at 37 °C in a humidified atmosphere containing 5% CO₂ and were confirmed to be free of mycoplasma contamination.

### Preparation of Photodegradable Gel Layers

Glass slides were coated by immersing in a type I collagen solution (Cellmatrix, Nitta Gelatin, Osaka, Japan) for at least 6 h, followed by rinsing the coated surface with Milli-Q water (Millipore Inc., Bedford, MA, USA) and air-drying. A total of 100 μL of 5% (w/v) gelatin from porcine skin (gel strength 300, type A) prepared in a 1:1 mixture of phosphate-buffered saline (PBS) and 0.3 M HEPES buffer (pH 7.0) was mixed with 100 μL of 2% (w/v) synthesized photocleavable crosslinker **2** solution prepared in 10 mM phthalic acid buffer containing 50 mM sodium chloride. The mixed solution was incubated in an aluminum block bath (ThermoUnit DTU-1B, TAITEC, Tokyo, Japan) preheated to 37 °C to initiate gelation. The partially gelled solution, prior to complete solidification, was applied onto the collagen-coated glass slides, and gel layers with defined thicknesses were formed using non-wire bar coaters of different sizes.

### PEG-Lipid-Modification and Light-Induced Cell Patterning

A bottomless microchannel (width: 3.8 mm, length: 18 mm, height: 0.40 mm; ibidi GmbH, Munich, Germany) was attached to the glass slide bearing the gel layer. A solution of 100 μM photocleavable PEG-lipid 1 prepared in a 1:1 mixture of PBS and DMSO was then introduced into the microchannel and incubated at 37 °C for 3 h to modify the gel surface. After washing with PBS, the microchannel was placed on a photomask, and light irradiation at 365 nm was applied using a photo-irradiator (REX-205, Asahi Spectra Co., Ltd., Tokyo, Japan) to activate the PEG-lipids. The released lipids were subsequently removed by washing with PBS. A cell suspension in PBS (1 × 10⁷ cells mL⁻¹) was then introduced into the microchannel and allowed to stand for 10 min to enable cell immobilization. Excess cells were removed by gentle washing with PBS. The patterned cells were observed using a fluorescence microscope (IX83, Olympus Co., Tokyo, Japan), and the acquired images were analyzed using ImageJ software. To prepare single-cell arrays, the gel surface was exposed to an array of light spots with varying diameters and two different inter-spot spacings. The number of cells per light-irradiated spot was determined by image analysis. To evaluate vertical positional changes of cells cultured on the gel layer, HT1080 cells stained with 25 μM CellTracker Green CMFDA (Thermo Fisher Scientific, C2925) were patterned on the gel surface and observed by confocal laser scanning microscopy (SP8, Leica Microsystems, Mannheim, Germany) before and after 14 h of incubation.

### Cell Release and Collection

Different procedures were employed for the retrieval of large numbers of cells and single cells. For bulk cell retrieval, a photo-irradiator was used to irradiate a large area of the gel surface. After irradiation, the microchannel was inverted and incubated at 37 °C for 1 h to facilitate gel degradation. PBS was then introduced from one side of the microchannel, and the released cells were collected from the opposite side. For single-cell retrieval, localized light irradiation was performed through the pinhole of a fluorescence microscope (10× objective, 5 s) to induce cell release. The released single cells were subsequently collected using a cell picker (Nepa Gene Co., Chiba, Japan) equipped with a 30 μm micropipette and transferred into either a 96-well plate or PCR tubes.

### Gene Expression Analysis

For RT–qPCR analysis, RNA from the collected cells was isolated according to the manufacturer’s instructions using a single-cell RNA purification kit (51800, Norgen Biotek Corp., Thorold, Canada). RT–qPCR was performed using a LightCycler 96 system (Roche Diagnostics K.K., Rotkreuz, Switzerland). The reactions were carried out with a One Step PrimeScript™ RT–PCR Kit (Perfect Real Time) (RR064A, Takara Bio Inc., Shiga, Japan) in accordance with the manufacturer’s protocol. The thermal cycling program consisted of a preincubation step at 42 °C for 30 s and 95 °C for 10 s, followed by 50 cycles of denaturation at 95 °C for 5 s and annealing/extension at 60 °C for 30 s. The primer sequences used in this study are listed in Table 1. For each target gene, expression levels were quantified relative to a control gene using the ΔΔCt method.

**Table 1.**
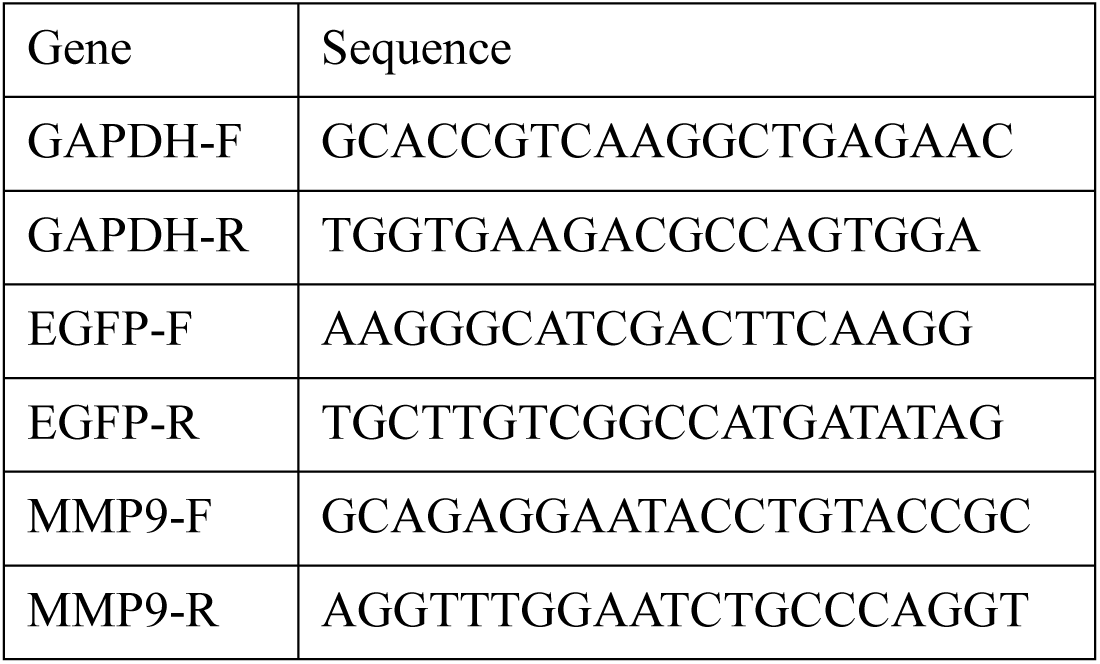
Primers for RT-qPCR.

### Single-Cell RNA-seq Analysis

For single-cell RNA sequencing (scRNA-seq) analysis, whole-transcriptome amplification of single-cell samples was performed using the Quartz-seq method according to the manufacturer’s protocol [54]. Briefly, single cells were lysed in lysis buffer and subsequently dissolved in priming buffer. mRNA from individual cells was denatured and primed by thermal cycling at 70 °C for 90 s followed by 35 °C for 15 s. Reverse transcription was then performed to synthesize first-strand cDNA by adding RT buffer, and SuperScript III (Thermo Fisher Scientific) followed by incubation at 35 °C for 5 min, 45 °C for 20 min, and 70 °C for 10 min. Subsequently, the excess primer was removed by treatment with Exo I buffer, with incubation at 37 °C for 30 min and enzyme inactivation at 80 °C for 20 min. The reaction mixture was then subjected to poly(A) tailing by adding poly(A) tailing buffer and incubating at 37 °C for 50 s, followed by heat inactivation at 65 °C for 10 min. PCR mix-1 was added, and the samples were incubated at 98 °C for 2 min 10 s, 40 °C for 1 min, and 68 °C for 5 min. Second-strand cDNA synthesis was carried out by adding PCR mix-2, followed by PCR amplification using the following cycling conditions: 21 cycles of 98 °C for 1 min, 65 °C for 15 s, and 68 °C for 5 min.

The amplified cDNA was purified using the MinElute PCR Purification Kit (QIAGEN, Tokyo, Japan). Sequencing libraries were prepared using the NEBNext Ultra RNA Library Prep Kit for Illumina (New England BioLabs, Massachusetts, USA) and sequenced on an Illumina HiSeq 2500 platform to generate 50-base single-end reads at a depth of 5 million reads per sample. Following sequencing, adapter sequences and low-quality reads were removed using Trimmomatic. The trimmed reads were aligned to the human reference genome (hg19) using HISAT2, and gene-level read counts were calculated with HTSeq-count. The resulting transcriptomic data were visualized and analyzed using R soft-ware.

### Single-Cell Image Analysis of Cellular Morphology

An array of single HT1080 cells on photodegradable gel layer was prepared as described above. The array was incubated in the DMEM supplemented with10 % FBS, 100 U/mL penicillin, and 100 μg/mL streptomycin at 37 °C in a humidified atmosphere containing 5% CO2 on the stage of a fluorescent microscope. Time-lapse images of the single-cell array were acquired every 5 min for 14 h and automatically analyzed as briefly described below (details are provided in the Supporting Information). All images were uniformly binarized to generate cell masks, and the centroid of each cell in the first frame was assigned to a 16 × 16 grid corresponding to one block of the single-cell array; each grid position was given a unique cell ID. Images were cropped into fixed regions centered on each grid position to generate single-cell image candidates. Valid single-cell images were identified using a Classification and Regression Tree (CART) model based on masking quality, cell multiplicity, and object length. The selected images were temporally linked to construct time-lapse sequences for each grid position.

For each cell, 13 morphological features were related to cell size, shape, and intensity-based texture were extracted from each frame. To summarize temporal behavior, the mean and standard deviation of each feature over the observation period were calculated, yielding a 26-dimensional morphological fingerprint for each cell. These morphological fingerprints were analyzed using UMAP to identify phenotypic clusters. Approximately 500 single-cell fingerprints obtained from prior validation experiments were used to construct a reference (pretrained) UMAP space. Single-cell data obtained from new experiments (target data) were processed using the same pipeline and overlaid onto the reference UMAP for cluster assignment. All analyses were implemented in a custom Python-based software system that outputs cluster identity, time-lapse image sequences, and spatial coordinates required for subsequent single-cell retrieval in real time.

## Acknowledgements

This work was supported by Japan Science and Technology Agency (JST) PRESTO (JPMJPR16FA), the MIRAI program (JPMJMI19G6) and CREST (JPMJCR2434). Part of this work was supported by research grants from the Uehara Memorial Foundation, the Ministry of Education, Culture, Sports, Science and Technology (MEXT) of Japan, Fund for Scientific Research (B) (24K01261), and JST A-STEP (JPMJTR254C). X.L. was supported by JST SPRING (JPMJSP2108).

## Data Availability Statement

The data that support the findings of this study are available from the corresponding author upon reasonable request.

