## Supplemental Figures for "Dual photoresponsive hydrogel surfaces enabling integrated single-cell analysis of morphology and gene expression"

### 1. Materials

4-[4-(1-Hydroxyethyl)-2-methoxy-5-nitrophenoxy]-butyric acid was purchased from Chemscene (State of New Jersey, USA). Diethyl ether, *N*-hydroxysuccinimide, *N*,*N'*-dicyclohexylcarbodiimide, and all anhydrous solvents were purchased from FUJIFILM Wako Pure Chemical Corporation (Osaka, Japan). Triethylamine was purchased from FUJIFILM Wako Pure Chemical Corporation and distilled before use. 1-ethyl-3-(3-dimethylaminopropyl) carbodiimide hydrochloride, Tris–HCl, and 4-nitrophenyl chloroformate were purchased from Tokyo Chemical Industry Co., Ltd. (Tokyo, Japan). Amine $PEG_4$ propionic acid and Sigmacote were purchased from Sigma-Aldrich Co. LLC (St. Louis, USA). All other solvents were purchased from Nacalai Tesque, Inc. (Kyoto, Japan). Pentaerythritol tetra(aminopropyl) polyoxyethylene (SUNBRIGHT PTE 100PA, Mw: 10,000) and pentaerythritol tetra(succinimidyloxyglutaryl) polyoxyethylene (SUNBRIGHT PTE 100GS, Mw: 10,000) were purchased from NOF Co. (Tokyo, Japan). Type I collagen solution (3.0 mg/mL, Cell Matrix Type I-A) was from Nitta Gelatin (Osaka, Japan). Sticky-slide VI 0.4 was from ibidi GmbH (Gräfelfing, Germany). Dulbecco's modified Eagles medium (DMEM) was from FUJIFILM Wako Pure Chemical Corporation. Penicillin-Streptomycin Mixed Solution and accutase were from Nacalai Tesque, Inc. Fetal bovine serum (FBS) was from Gibco (Thermo Fisher Scientific, Carlsbad, CA, USA) and Nichirei Biosciences Inc. (Tokyo, Japan). CellTracker Green CMFDA and 5-TAMRA were from Thermo Fisher Scientific (Waltham, MA, USA). Phalloidin-iFluor 555 Reagent and Alexa Fluor® 647 Anti-Vinculin antibody were purchased from Abcam plc, (Cambridge, UK). All other reagents were of analytical grade.

### 2. General procedures

$^1$H-NMR spectra were recorded on JEOL NMR JNM-ECZ600R (JEOL Ltd., Tokyo, Japan), and all chemical shifts are reported in ppm downfield of tetramethylsilane using residual solvent as internal reference. Light irradiation was performed with an ultraviolet (UV) irradiator (REX-250, from Asahi Spectra Co., Ltd., Tokyo, Japan) equipped with a cylindrical lens through a bandpass filter (wavelength: 365 ± 5 nm). Photomasks were purchased from Nippon Filcon (Tokyo, Japan). Microscopy was performed by inverted fluorescence microscope (Cat#: IX83, Olympus Corporation, Tokyo, Japan) with Stage Top Incubator (Cat#: STXG-IX3WX-SET, Tokai Hit., Co, Ltd., Shizuoka, Japan). Confocal microscopy was performed by Leica TCS SP8 confocal microscope (Leica Microsystems, Wetzlar, Germany). Cell picking was finished by a capillary-based cell picker (NEPA GENE, Chiba, Japan). All solvents and reagents were commercially available and used without further purification.

### 3. Synthesis of photocleavable crosslinker **2**

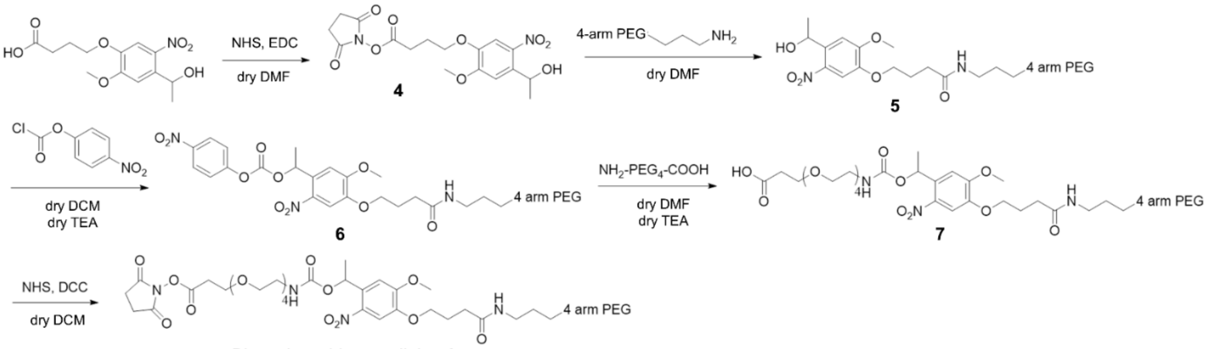


**Figure S1 | Synthetic scheme of photocleavable crosslinker 2.**

#### Synthesis of compound **4**

4-[4-(1-Hydroxyethyl)-2-methoxy-5-nitrophenoxy]-butyric acid (321.9 mg, 1.07 mmol, 1 eq), *N*-hydroxysuccinimide (247.6 mg, 2.15 mmol, 2 eq) and 1-ethyl-3-(3-dimethylaminopropyl) carbodiimide hydrochloride (206.2 mg, 1.07 mmol, 1 eq) were dissolved in dry dimethylformamide (3 mL) and stirred in dark for 24 hours under room temperature. After TLC monitoring, the solvent was removed under reduced pressure. Water (10 mL) was added to the residue, and the precipitated solid was collected by vacuum filtration and dried to give the desired product as a yellow solid (380.7 mg, >99.9%) : **$^1$H-NMR** (600 MHz, DMSO-*d6*) δ 1.36-1.37 (d, *J* = 6.0 Hz, 3H, $CHCH_3$), 2.06-2.12 (m, 2H, $CH_2CH_2CH_2$), 2.82 (s, 4H, $COCH_2CH_2CO$), 2.85-2.88 (t, *J* = 6.0 Hz, 2H, $COCH_2CH_2$), 3.91 (s, 3H, $OCH_3$), 4.12-4.14 (t, *J* = 6.0 Hz, 2H, $CH_2CH_2O$), 5.24-5.29 (m, 1H, $CH_{Ar}CHOH$), 7.37 (s, 1H, $CH_{Ar}$), 7.55 (s, 1H, $CH_{Ar}$).

#### Synthesis of compound **5**

Compound **4** (55.2 mg, 0.14 mmol, 8 eq) and pentaerythritol tetra(aminopropyl) polyoxyethylene ($M_W$: 10000) (180.0 mg, 0.018 mmol, 1 eq) were dissolved by 10 mL dry dimethylformamide and stirred in dark for 24 hours under room temperature. After TLC monitoring, the solvent was removed under reduced pressure. Tris–HCl buffer (10 mL, pH 8.0) was added to the residue to quench the reaction, and the mixture was centrifuged (12,000 × g, 4 °C, 10 min). The clear liquid was transferred to a dialysis tube (MWCO 3.5 kDa) and dialyzed against water for 24 h. The solution inside the tube was then freeze-dried to give the product as a light yellow solid (195.8 mg, 97.7%) : **$^1$H-NMR** (600 MHz, DMSO-*d6*) δ 1.36-1.37 (d, *J* = 6.0 Hz, 3H, $CHCH_3$), 1.57-1.64 (m, 2H, $NHCH_2CH_2$), 1.91-1.97 (m, 2H, $CH_2CH_2CO$), 2.21-

2.24 (t, $J = 6$ Hz, 2H, C$H_2$CONH), 3.06-3.10 (quart., $J = 12.0$ Hz, 2H, C$H_2$($CH_2O$)$_n$), 3.32-3.55 (brs, ($CH_2O$)$_n$), 3.91 (s, 3H, $OCH_3$), 3.64-3.66 (t, $J = 6.0$ Hz, 2H, NHC$H_2$$CH_2$), 3.90 (s, 3H, $OCH_3$), 4.01-4.04 (t, $J = 6.0$ Hz, 2H, OC$H_2$$CH_2$), 5.24-5.27 (m, 1H, HOC$H$$CH_{Ar}$), 5.46-5.47 (d, $J = 6.0$ Hz, 1H, CON$H$), 7.36 (s, 1H, $CH_{Ar}$), 7.51 (s, 1H, $CH_{Ar}$), 7.81-7.83 (t, $J = 6.0$ Hz, 1H, $H$O$CH_2$).

Synthesis of compound **6**

Compound **5** (153.7 mg, 0.014 mmol, 1 eq), 4-nitrophenyl chloroformate (127.8 mg, 0.634 mmol, 45 eq), dry dichloromethane (10 mL) and dry triethylamine (0.1 mL) were mixed and stirred in dark for 3 days under room temperature. After TLC monitoring, the solvent was removed under reduced pressure. The residue was dissolved again in dichloromethane (2.5 mL) and added dropwise to cold diethyl ether (40 mL). The precipitate recovered by centrifugation (12,000 × g, 4 °C, 10 min) and air-dried to give the product as a pale flesh-colored solid (260.4 mg, >99.9%) : **$^1$H-NMR** (600 MHz, DMSO-*d6*) δ 1.58-1.64 (m, 2H, NH$CH_2$C$H_2$), 1.72-1.73 (d, $J = 6.0$ Hz, 3H, CHC$H_3$), 1.94-1.97 (m, 2H, $CH_2$C$H_2$CO), 2.22-2.25 (t, $J = 6.0$ Hz, 2H, C$H_2$CONH), 3.28-3.55 (brs, ($CH_2O$)$_n$), 3.64-3.65 (t, $J = 6.0$ Hz, 2H, NHC$H_2$$CH_2$), 3.96 (s, 3H, $OCH_3$), 4.05-4.08 (t, $J = 6.0$ Hz, 2H, OC$H_2$$CH_2$), 6.26-6.29 (m, 1H, OC$H$$CH_{Ar}$), 7.19 (s, 1H, $CH_{Ar}$), 7.52-7.54 (d, $J = 6.0$ Hz, 2H, C$H_{Ar}$O), 7.51 (s, 1H, $CH_{Ar}$), 8.29-8.30 (d, $J = 6.0$ Hz, 2H, $O_2$NC$H_{Ar}$).

Synthesis of compound **7**

Compound **6** (164.1 mg, 0.014 mmol, 1 eq), amine $PEG_4$ propionic acid (55.7 mg, 0.210 mmol, 15 eq), dry dimethylformamide (10 mL) and dry triethylamine (0.1 mL) were mixed and stirred in dark for 24 hours under room temperature. After TLC monitoring, the solvent was removed under reduced pressure. Water (15 mL) was added to the residue and the mixture was centrifuged (12,000 × g, 4 °C, 10 min). The clear liquid was transferred to a dialysis tube (MWCO 3.5 kDa) and dialyzed for 24 h. The solution inside the tube was then freeze-dried to give the product as a white solid (134.4 mg, 78.5%) : **$^1$H-NMR** (600 MHz, DMSO-*d6*) δ 1.51-1.52 (d, $J = 6.0$ Hz, 3H, CHC$H_3$), 1.59-1.64 (m, 2H, NH$CH_2$C$H_2$), 1.93-1.97 (m, 2H, $CH_2$C$H_2$CO), 2.21-2.24 (t, $J = 6.0$ Hz, 2H, C$H_2$CONH), 2.42-2.44 (t, $J = 6.0$ Hz, 2H, C$H_2$COOH), 3.07-3.12 (m, 4H, C$H_2$($CH_2O$)$_n$, ($CH_2O$)$_n$C$H_2$), 3.28-3.60 (brs, ($CH_2O$)$_n$), 3.64-3.65 (t, $J = 6.0$ Hz, 2H, NHC$H_2$$CH_2$), 3.91 (s, 3H, $OCH_3$), 4.03-4.06 (t, $J = 6.0$ Hz, 2H, $CH_{Ar}$OC$H_2$), 6.09-6.13 (m, 1H, OC$H$$CH_{Ar}$), 7.13 (s, 1H, $CH_{Ar}$), 7.39-7.41 (t, $J = 6.0$ Hz, 1H, $CH_{Ar}$N$H$CO), 7.56 (s, 1H, $CH_{Ar}$), 7.81-7.83 (t, $J = 6.0$ Hz, 1H, CON$H$$CH_2$).

Synthesis of photocleavable crosslinker **2**

Compound **7** (91.2 mg, 0.007 mmol, 1 eq), *N*-hydroxysuccinimide (21.9 mg, 0.190 mmol, 27 eq) and *N*,*N*'-dicyclohexylcarbodiimide (62.8 mg, 0.304 mmol, 43 eq) were dissolved in dry dichloromethane (10 mL) and stirred in dark for 24 hours under room temperature. The residue was dissolved again in dichloromethane (3 mL) and added dropwise to cold diethyl ether (40 mL). The precipitate recovered by centrifugation (12,000 × g, 4 °C, 10 min) and air-dried. The purification was done twice to give the product as a pale flesh-colored solid (75.2 mg, 80.0%) : $^1$**H-NMR** (600 MHz, $CDCl_3$) δ 1.58-1.59 (d, *J* = 6.0 Hz, 3H, $\mathrm{CHC}H_3$), 1.75-1.80 (m, 2H, $\mathrm{NHCH_2C}H_2$), 2.17-2.20 (m, 2H, $\mathrm{CH_2C}H_2\mathrm{CO}$), 2.35-2.38 (t, *J* = 9.0 Hz, 2H, $\mathrm{C}H_2\mathrm{CONH}$), 2.82 (s, 4H, $\mathrm{COC}H_2\mathrm{C}H_2\mathrm{CO}$), 2.87-2.93 (m, 2H, $\mathrm{C}H_2\mathrm{COO}$), 3.31-3.69 (brs, $(CH_2O)_n$), 3.77-3.79 (t, *J* = 6.0 Hz, 2H, $\mathrm{COC}H_2\mathrm{CH_2}$), 3.83-3.86 (t, *J* = 6.0 Hz, 2H, $\mathrm{NHC}H_2\mathrm{CH_2}$), 3.95 (s, 3H, $OCH_3$), 4.08-4.11 (t, *J* = 6.0 Hz, 2H, $\mathrm{CH_{Ar}OC}H_2$), 5.40 (s, 1H, $\mathrm{CH_{Ar}N}H\mathrm{CO}$), 6.35-6.36 (d, *J* = 6.0 Hz, 2H, $\mathrm{CON}H\mathrm{CH_2}$, $\mathrm{OC}H\mathrm{CH_{Ar}}$), 7.01 (s, 1H, $CH_{Ar}$), 7.58 (s, 1H, $CH_{Ar}$).

### 4. Microchannel system

A commercially available bottomless six-channel coverslip (Sticky-Slide VI 0.4; Cat. No. 80608, ibidi GmbH, Gräfelfing, Germany) was used for cell culture experiments. The coverslip was directly adhered to the hydrogel layer to form a microchannel system and secured using a custom-made fixture (Figure S2).

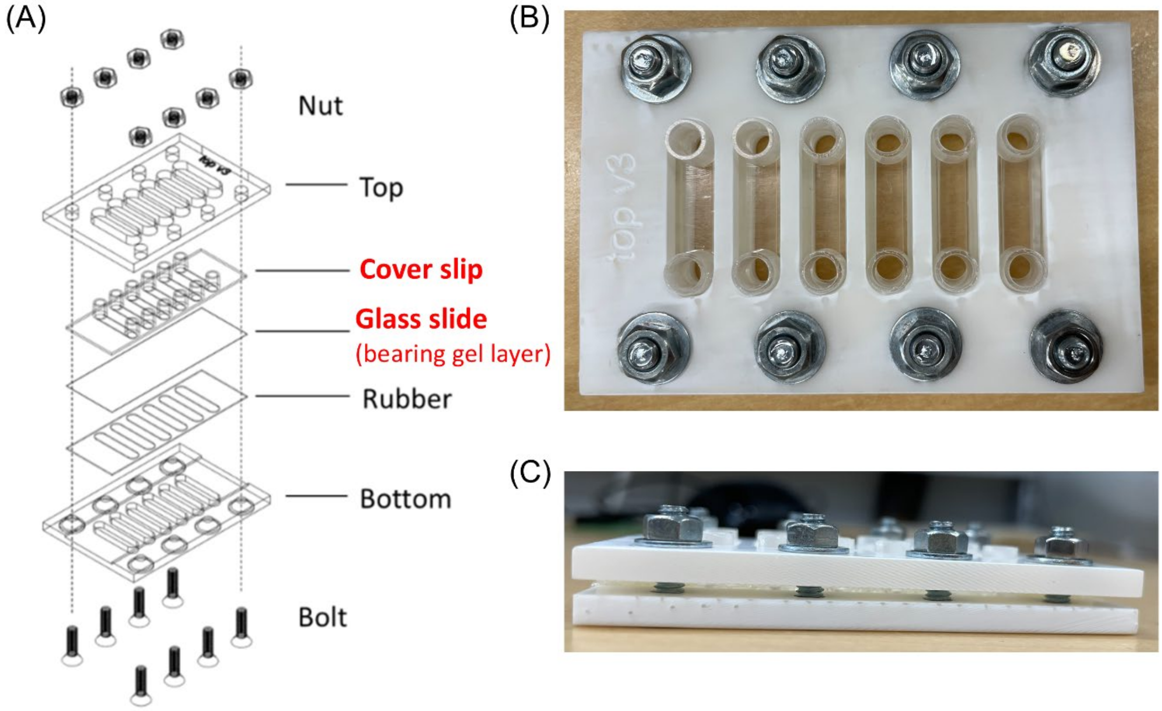

**Figure S2 | Design drawings and photographs of the tapeless microchannel system.** (A) Layout design of the system. A fixture jig fabricated by 3D printing was used to ensure tight contact between the coverslip and the glass slide. (B) Top-view photograph of the system. (C) Side-view photograph of the system.

### 5. Gel degradation test

The photodegradable gel layer modified with PEG-lipid **3** was evaluated for its ability to undergo light-induced degradation and cell release upon irradiation at 365 nm. Various irradiation doses were applied to determine the minimum light exposure required for gel degradation and effective cell release (Figure S3).

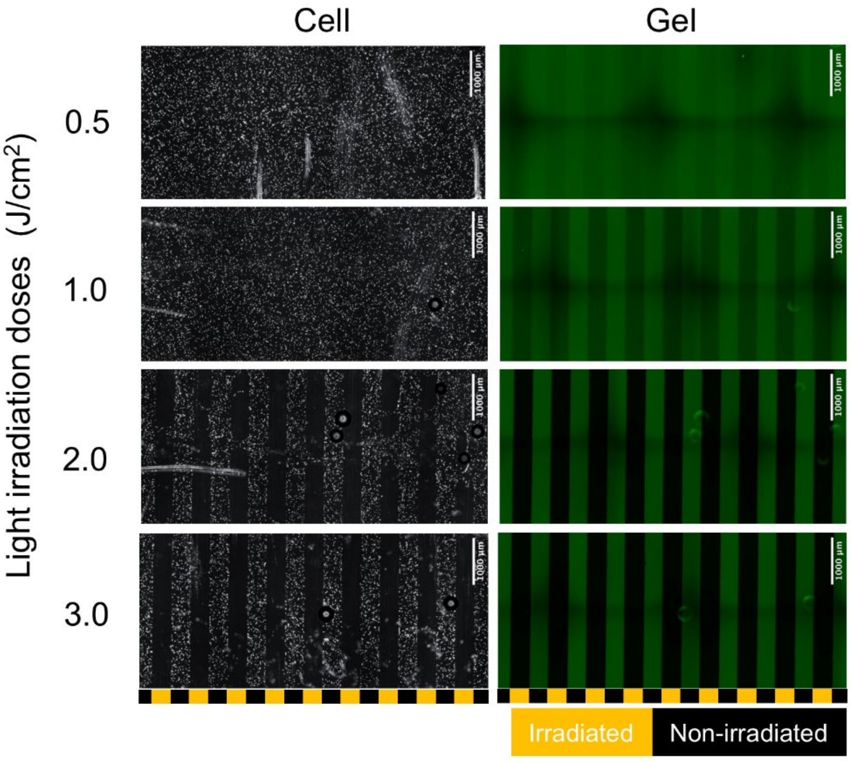


**Figure S3 | Fluorescence microscopy images of anchored cells on green fluorescence–labeled gel layers after exposure to a line-patterned light at varying irradiation doses.** The gel layers were labeled with 100 mM fluorescein-5(6)-carboxamidocaproic acid N-hydroxysuccinimidyl ester. (Left column) Bright-field images of HeLa cells. (Right column) Green fluorescence images of the gel layers. Scale bar: 1000 μm.

### 6. Light-induced cell patterning

To examine whether cells can be photo-patterned on a 4-arm PEG–crosslinked gelatin gel layer by modification with photoactivatable PEG-lipid **1**, a non-photodegradable gelatin gel layer crosslinked with non-photocleavable 4-arm PEG was modified with PEG-lipid **1** and exposed to patterned light irradiation at 365 nm with varying doses. After seeding HeLa cells and rinsing, the density of anchored cells on the gel layers increased proportionally with the irradiation dose, indicating dose-dependent control of cell attachment (Figure S4).

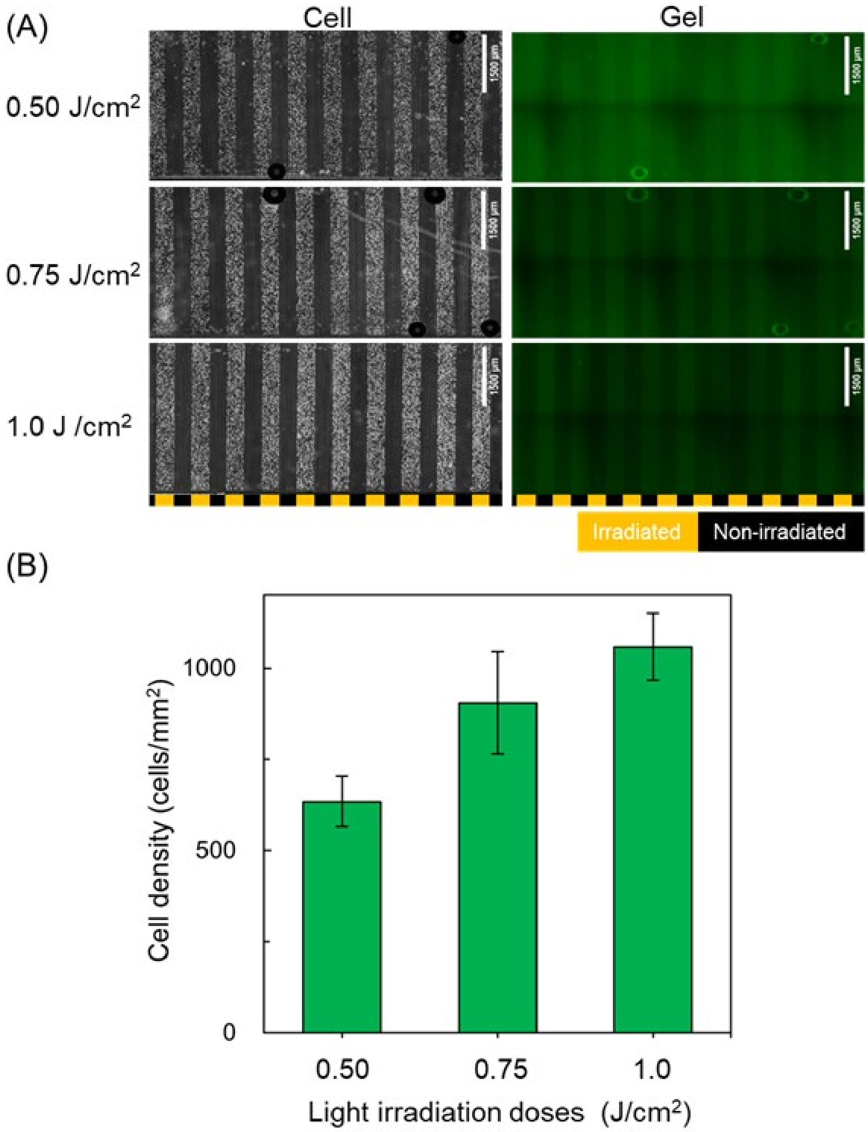


**Figure S4 | Light-induced cell anchoring on a gelatin-based gel layer modified with photoactivatable PEG-lipid.** (A) Fluorescence microscopy images of anchored HeLa cells on fluorescein-stained gel layers after exposure to line-patterned light at varying irradiation doses. Scale bar: 1000 μm. (B) Quantified cell densities at different irradiation doses.

### 7. Cell adhesion onto gel layers

To examine whether adherent cells can properly attach to gel-layer surfaces modified with photoactivatable PEG-lipid 1, anchored HeLa cells were stained with a fluorescently labeled anti-vinculin antibody (Alexa Fluor 647–labeled anti-vinculin antibody) and an actin-staining reagent (phalloidin–iFluor 555) after 11 h of incubation. Non-photodegradable gel layers were prepared by crosslinking gelatin with non-photocleavable 4-arm PEG functionalized with NHS esters at each terminus. The gel layers were modified with 20 μM photoactivatable PEG-lipid 1 and exposed to an array of light spots. HeLa cells were then seeded onto the surface, and after a 10 min incubation, the surface was rinsed with PBS to form single-cell arrays. As a control, HeLa cells were randomly seeded onto gel layers without PEG-lipid modification. Both surfaces were incubated under standard cell culture conditions, after which the cells were fixed and permeabilized, followed by fluorescent staining. As shown in Figure S6, actin filaments and focal adhesions involving vinculin were successfully stained in red and green, respectively. No significant differences were observed between the two conditions, indicating that cell adhesion to the gel-layer surface was not adversely affected by PEG-lipid modification.

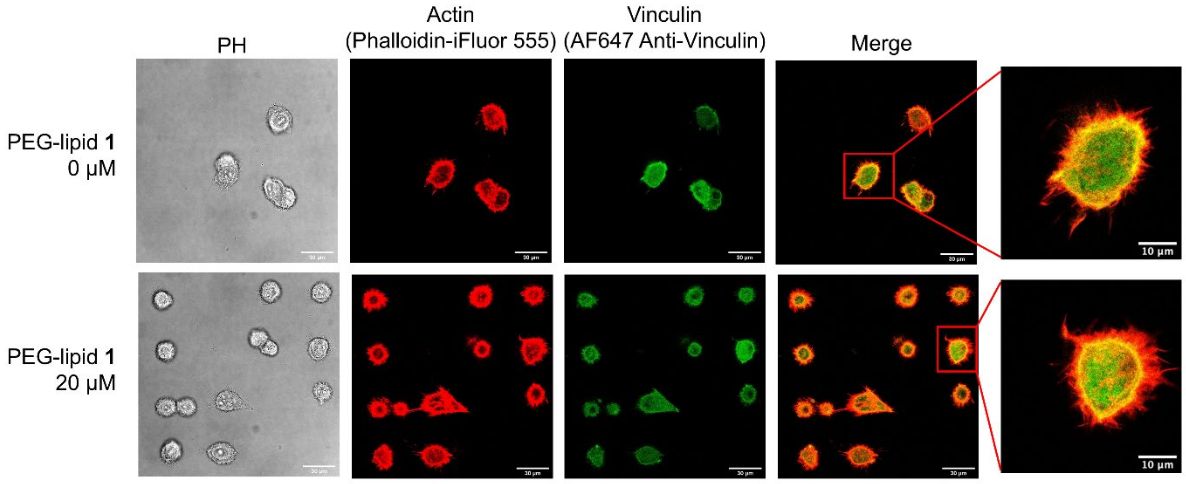


**Figure S5 | Fluorescence microscopy images of stained HeLa cells after 11 h of incubation on gel layers without or with photoactivatable PEG-lipid 1.** (*Upper*) HeLa cells cultured on gel layers without PEG-lipid modification. (*Lower*) HeLa cells arrayed and cultured on gel layers modified with 20 μM PEG-lipid **1**. Actin filaments are stained with fluorescently labeled phalloidin in red, and focal adhesions are stained with fluorescently labeled anti-vinculin antibody in green. Scale bars: 30 μm and 10 μm.

### 8. Light-induced cell retrieval and gene expression analysis

We examined whether cells retrieved from the gel-layer surface could be analyzed by RT–qPCR. HeLa cells and HeLa–EGFP cells were independently anchored in line patterns on dual photoresponsive gel layers that were fluorescently stained with rhodamine. Entire, half, or quarter regions of the cell-patterned surfaces were exposed to light at 365 nm, after which the microchannel was inverted and incubated at 37 °C for 1 h. PBS was then introduced from the inlet of the microchannel, and the released cells were collected by pipetting from the outlet on the opposite side of the microchannel.

As shown in Figure S6A, both the anchored cells and the red-fluorescent gel layers disappeared from the light-irradiated regions after PBS introduction and pipetting. The collected cell suspensions were transferred into PCR tubes, and RNA was isolated using a commercially available RNA purification kit. The EGFP transcript levels were subsequently analyzed by RT–qPCR. As shown in Figure S6B, EGFP transcripts were detected exclusively in the recovered HeLa–EGFP cells, and gene expression levels could be quantitatively evaluated in proportion to the light-irradiated area.

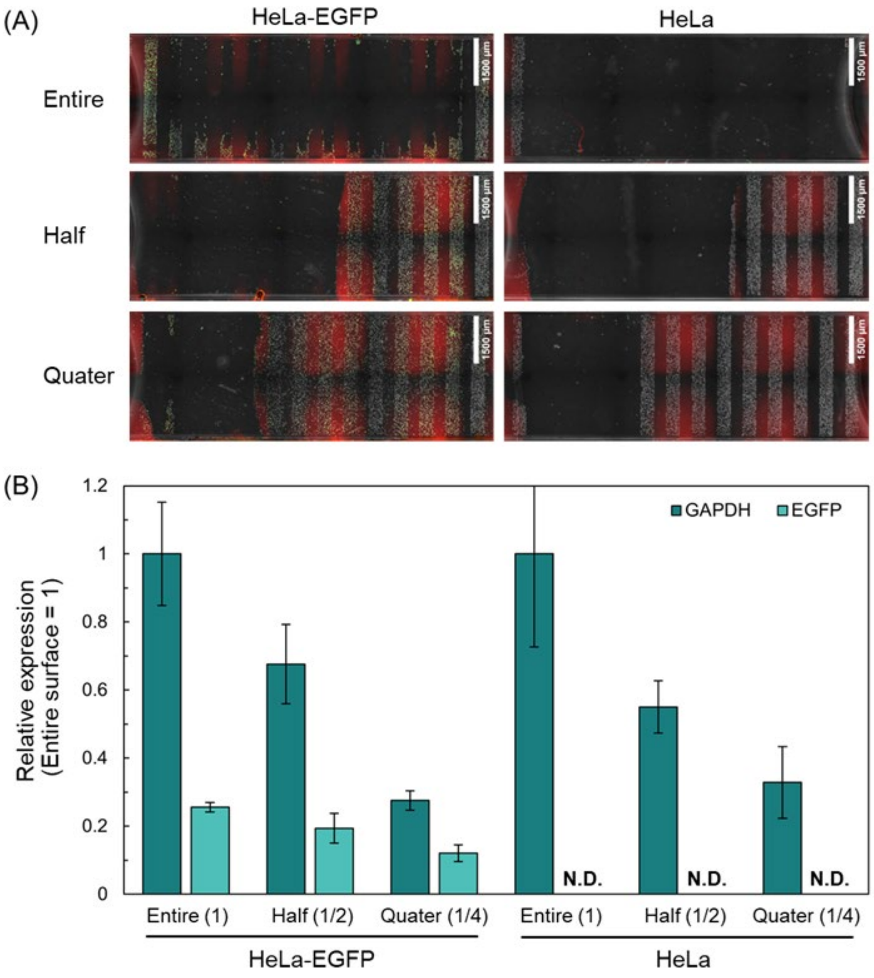

**Figure S6 | Light-induced cell retrieval and gene expression analysis.** (A) Fluorescence microscopy images of (left) HeLa cells and (right) HeLa–EGFP cells on rhodamine-stained gel layers after light irradiation of entire, half, or quarter regions of the gel surface. Scale bar: 1500 μm. (B) Relative expression levels of GAPDH and EGFP in the collected cells.

### 9. Automated time-course morphological profiling

All images acquired from the single-cell array were processed using a uniform binarization threshold based on Otsu's method to generate cell masks. Using the first frame of the time-lapse sequence, the centroid of each masked cell was calculated to determine its initial seeding position. Centroid coordinates were classified into 16 bins along both the x- and y-axes, corresponding to the 16 × 16 grid structure of the single-cell array. Each cell was associated with the corresponding grid intersection coordinate and given a unique cell ID. Based on these grid coordinates, images were cropped into fixed regions (50 × 50 pixels) centered on each grid position to generate cropped images containing individual cells. To identify valid single-cell images, the cropped images were classified using a CART (Classification and Regression Tree) model trained on approximately 256 manually annotated single-cell images. The model categorizes images into four classes based on masking quality, cell multiplicity within the cropped region, and object length (≤ or > 12 pixels). This CART model achieved 100% classification accuracy on the labeled dataset. Only cropped images classified as high-confidence single-cell images ("single cell-defined grid images") were retained. These images were temporally linked to construct time-lapse image sequences for each grid position.

For each time-lapse sequence, 13 morphological features were extracted from a single cell in each frame: area, length, width, length–width ratio, perimeter, compactness, mean intensity, intensity standard deviation, total intensity, correlation, energy, homogeneity, and contrast. This feature set was selected based on prior validation in our group, in which these variables showed relatively low redundancy while enabling successful machine-learning–based prediction of multiple cell quality attributes.

To summarize temporal behavior, the mean and standard deviation of each feature were calculated over the observation period, resulting in a 26-dimensional morphological fingerprint (13 features × 2 statistics) for each single cell. In this study, fingerprints were derived from 14 hours of time-lapse imaging. Cells were excluded from analysis if the time series showed cell loss, cell entry from neighboring grids, or other disruptions, even if the grid was initially classified as containing a single cell. As a result, only cells with complete and stable 14-hour fingerprints were included in downstream analyses.

Using these morphological fingerprints, UMAP analysis was performed to identify distinct morphological clusters. Approximately 500 single-cell fingerprints obtained from prior validation experiments were used to define the reference UMAP space; these data are referred to as pre-trained data in this study. After cluster structures were established and labeled using the pre-trained data, single-cell data acquired from newly performed experiments were processed using the same image analysis pipeline. These newly acquired data were treated separately as target data. Following real-time analysis, the target data were overlaid with the pre-trained data to visualize cluster assignment for each target cell. This workflow was implemented as a custom Python-based software system that automatically outputs cluster identity, time-lapse image sequences, and spatial coordinates required for physical cell collection. All analysis and outputs were performed in real time in single cell collection experiment.

### 10. NMR spectra of new compounds

Compound **5**

$^{1}$H-NMR

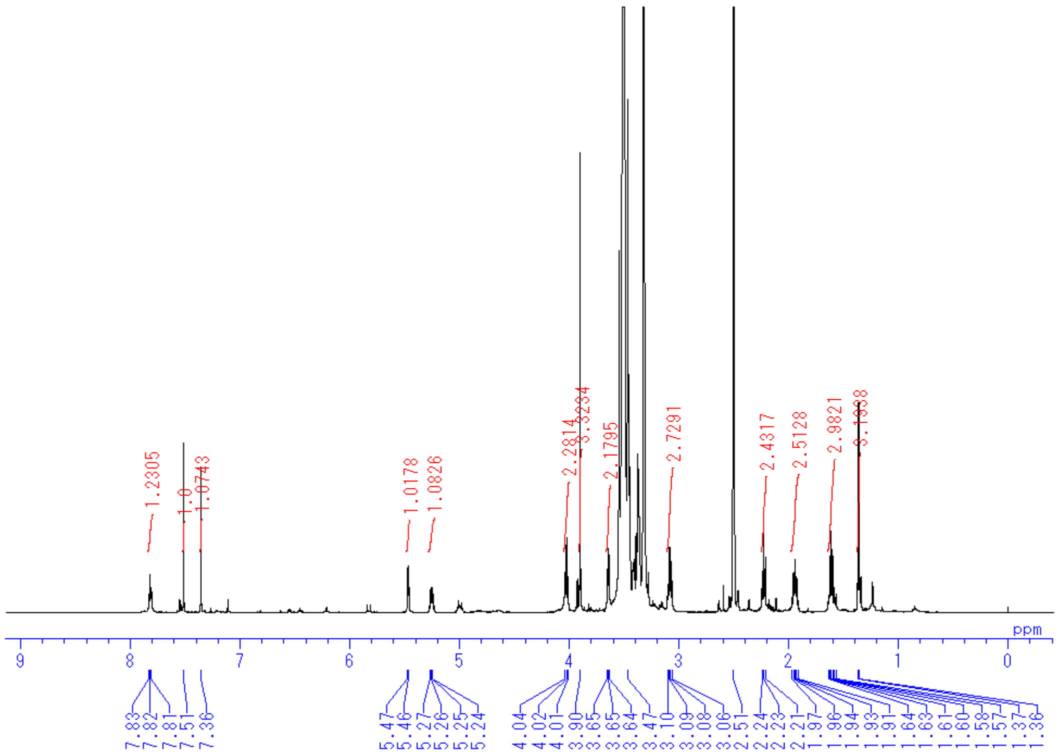

Compound **6**

$^{1}$H-NMR

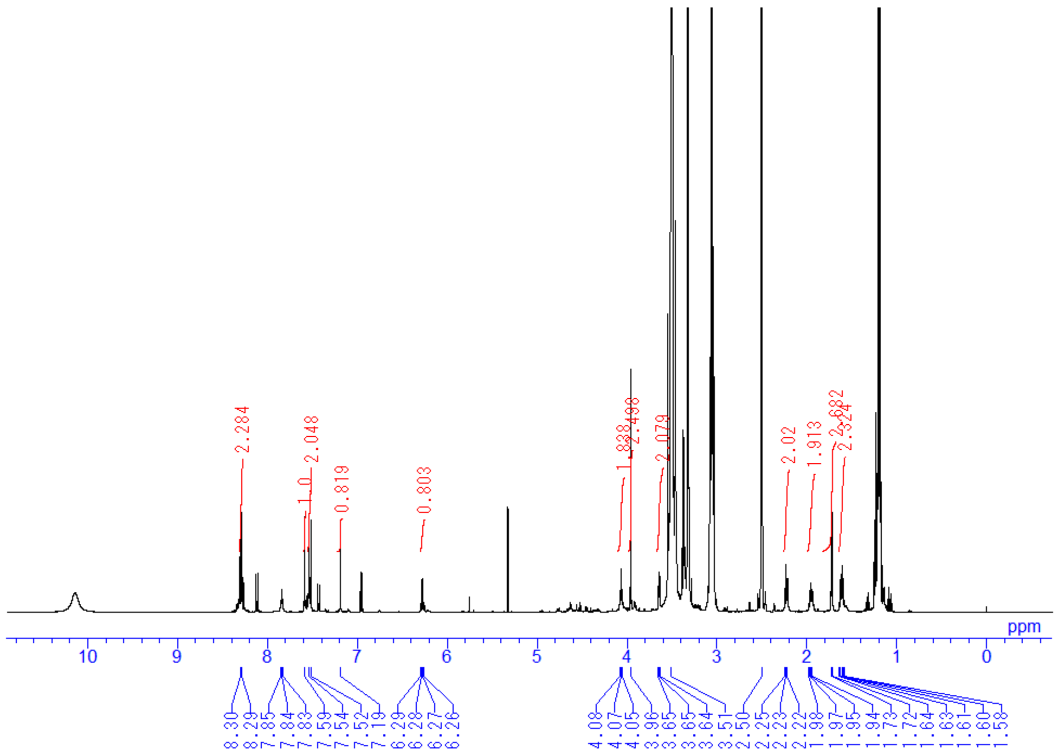

Compound **7**

$^1$H-NMR

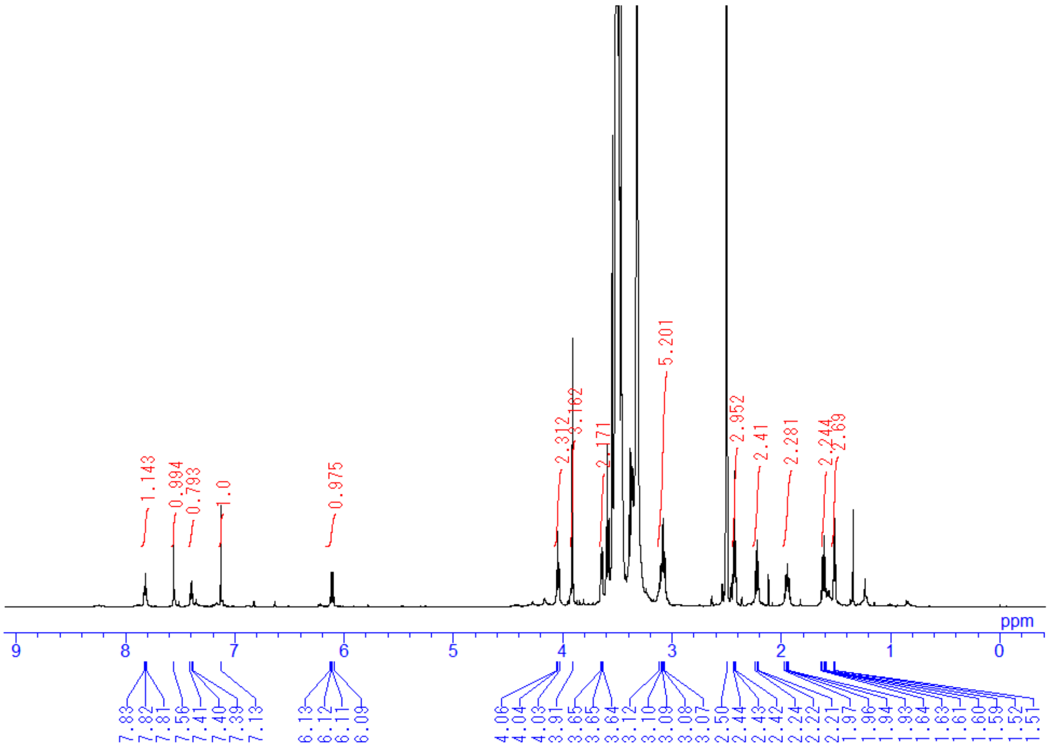


Photocleavable crosslinker **2**

$^1$H-NMR

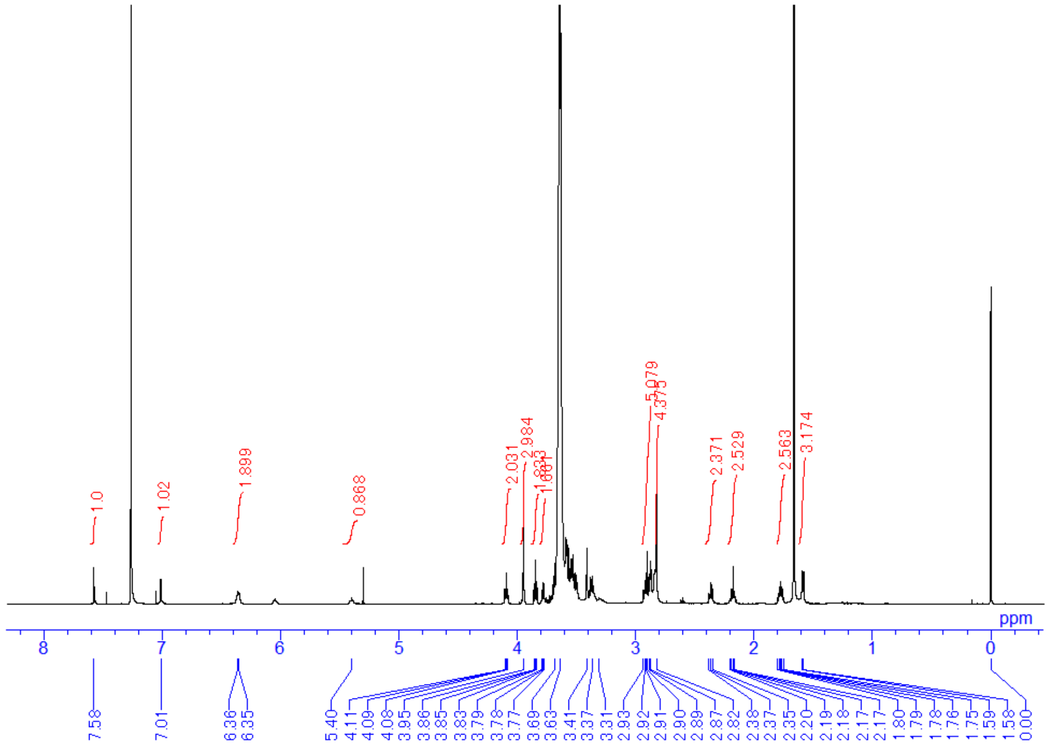
